# Molecular basis of noncanonical complement C3 activation by histamine

**DOI:** 10.64898/2025.12.13.694092

**Authors:** Francisco J. Fernández, Karla de la Paz-García, Javier Querol-García, Carlos A. Ramos-Guzmán, Héctor Martín-Merinero, Israel Mares-Mejía, Lucía Alfonso-González, Santiago Rodríguez de Córdoba, Iñaki Tuñón, M. Cristina Vega

## Abstract

For fifty years the tick-over mechanism has been considered responsible for priming the activation of the complement system’s alternative pathway through the reaction of a nucleophilic water molecule with C3 yielding C3(H_2_O), even though the exclusivity of this role has been challenged by the existence of extrinsic proteases that can cleave circulating C3 into C3b. Here we show that the biogenic amine histamine can activate C3 by reacting with the internal thioester bond yielding a novel species that is equivalent to C3(H_2_O), which we have called C3h. Histamine activation of C3 occurs significantly faster than the water-mediated tick-over reaction, leading to the accelerated release of the C3a anaphylatoxin moiety, contributing to inflammation. Importantly, C3h can form an active C3 convertase enzyme that, together with the released C3a, can amplify complement activation and inflammatory responses. These results offer insight into the priming of the complement system activation and support the existence of direct crosstalk between histamine-releasing processes and complement activation.

## Introduction

The complement system of the mammalian immune system is a protein network that contains zymogens, enzymes, soluble and membrane-associated receptors and regulators, which collectively protect the organism against bacterial, viral, and eukaryotic pathogens, and help to maintain homeostasis by removing immune complexes and clearing abnormal cells and cellular debris (*1*–*3*). C3, the central complement component, circulates as an inert protein until it becomes proteolytically activated by multiprotein enzymes called C3 convertases (*4*). These enzymes contain an active form derived from C3, C3b, which explains the self-amplifying nature of the alternative pathway. Paradoxically, C3 convertases require a small amount of activated C3 to prime them, otherwise they cannot assemble properly. The tick-over mechanism was proposed to resolve this paradox by invoking the existence of a low level of an activated species derived from C3 by reaction of the internal high-energy isoglutamyl cysteine thioester bond (Cys1010-Gln1013) with the ubiquitous water nucleophile, which results in an acyl-imidazole intermediary that can react preferentially with hydroxyl nucleophiles (*5*). The breakdown of the thioester bond in fluid phase triggers a slow conformational change that results in C3 adopting a conformation most closely resembling C3b, termed C3(H_2_O). In addition, extrinsic proteases can proteolytically activate C3 by cleaving off C3a (*6*–*8*). Furthermore, C3 can also be regarded as a contact activated protein (*9*), as it can be activated by interaction with a variety of biological surfaces, apparently by-passing the need for a first-hit reaction with water or the action of nonspecific proteases.

In contrast to C3b, which can react with hydroxyl nucleophiles on the cell surface and become covalently attached to surfaces, C3(H_2_O) remains in the fluid phase. Before it is removed by proteolytic cleavage by complement regulators like factor H (FH) and factor I (FI), C3(H_2_O) can assemble catalytically proficient C3 convertases and turn over C3 into C3b. Due to the extremely low quantities of C3(H_2_O) that exist in blood and the relatively rapid turnover of C3(H_2_O) into its inactivated fragments, the tick-over mechanism provides an explanation for the seemingly contradictory necessity of a primed alternative pathway. As the reaction with water is slow and inefficient, small primary amines like ethanolamine or methylamine have been used to chemically produce analogous C3(H_2_O) species for their characterization. However, no biological role has been attributed to these activated species as the primary amines used to activate C3 are not naturally present in the organism.

Histamine is a biogenic amine with a vast array of biological activities (*10*, *11*). It has been shown to play a key physiological role in the control of gastric acid secretion and a pathophysiological role in a range of allergic disorders (*10*). Histamine is found at low concentrations in the blood of healthy individuals, with concentrations ranging between 2.7–9.0 nM in plasma (*12*–*14*) and 225–585 nM in blood (*15*), and at much greater concentrations in certain pathological states like anaphylaxis (50–300 nM in plasma), septic shock (200–500 nM in plasma), and acute coronary disease (50–150 nM in plasma, 810–1140 nM in blood) (*15*). Histamine can reach higher concentrations locally during degranulation of mast cells and basophils (estimated at 0.1–1 mM) (*16*). One of the best-known functions of histamine is as a mediator of the potent inflammatory response mounted during allergy. Upon contact with the allergic stimulus, basophils and mast cells release histamine and other biologically active molecules that enhance vascular permeability and stimulate a local inflammatory state. Research has established that histamine can modulate the release of various proinflammatory interleukins and cytokines, which also help control inflammation (*17*). When the allergic stimulus is not properly controlled and downregulated, excessive inflammation can cause cell and tissue damage.

The role of the complement system in promoting histamine release and other responses associated with allergy and autoimmunity has been demonstrated previously (*18*). Specifically, the anaphylatoxins C3a and, especially, C5a have been shown to stimulate histamine release through the direct binding to their cognate receptors in basophils and mast cells (*19*) or by stimulating the metabolism of arachidonic acid (*18*). In addition, C3a could play additional roles in hypersensitivity responses beyond inducing histamine release, e.g., by recruiting basophils, which express C3aR, to inflammation sites (*18*). This is particularly interesting as C3a has been shown to be able to form through the proteolytic activity of FXa and FIIa during the activation of the coagulation extrinsic pathway (*18*).

However, the potential roles of released histamine in modulating the complement system remain largely uncharacterized and poorly understood. Early work reported that histamine has either an inhibitory or an enhancing effect on the biosynthesis of complement factors by monocytes (*20*) and hepatoma cells (*21*), depending on the histamine receptor that mediated the effect, with the H_1_ receptor mediating the increase and the H_2_ receptor the decrease in complement factor synthesis and secretion (*21*). Beyond this nonspecific effect on complement factor biosynthesis, no direct reaction or interaction between histamine and complement factors has yet been reported.

In this work, we describe a novel form of C3 formed by the reaction of histamine with the thioester bond in the TED domain of C3. This species, denominated C3h, is activated by a conformational change triggered by reaction with histamine, which we show to resemble most closely the structure of C3b by X-ray crystallography. In agreement with its C3b-like structure, C3h can assemble a functional C3 convertase, activating the alternative complement pathway. Additionally, C3h is rapidly converted into C3b-histamine (C3bh) and C3a by spontaneous chemical cleavage and proteolyzed into C3dg and C3c by the action of FI. Therefore, C3h bears all the hallmarks of C3(H_2_O). We propose that C3h may have a proinflammatory role under conditions characterized by high acute or chronic histamine concentration, like allergic reactions, autoimmunity, and chronic cardiovascular disease, amplifying the strong inflammatory stimulus that histamine release provides through complement activation.

## Results

### Histamine forms a covalent adduct with C3

To determine whether the biogenic amine histamine reacts with the thioester bond of C3 (Cys1010-Gln1013), we incubated C3 purified from human blood at the physiological concentration of 1 mg/ml (∼5.4 µM) with 100 mM histamine in phosphate buffered saline (pH 7.4) at 4 °C for 12 h (**Fig. 1A,B**). Then we buffer exchanged the reaction mixture to remove excess histamine and separated reacted C3 from unreacted C3 by anion exchange chromatography (**Fig. 1C**). Tellingly, the elution volume of histamine-reacted C3 was nearer the expected elution volume of C3(H_2_O) than to that of C3, therefore allowing a complete resolution of the two species (*22*, *23*). We refer to the resultant histamine-adduct of C3 as C3h (“C3 histamine”).

**Fig. 1.**
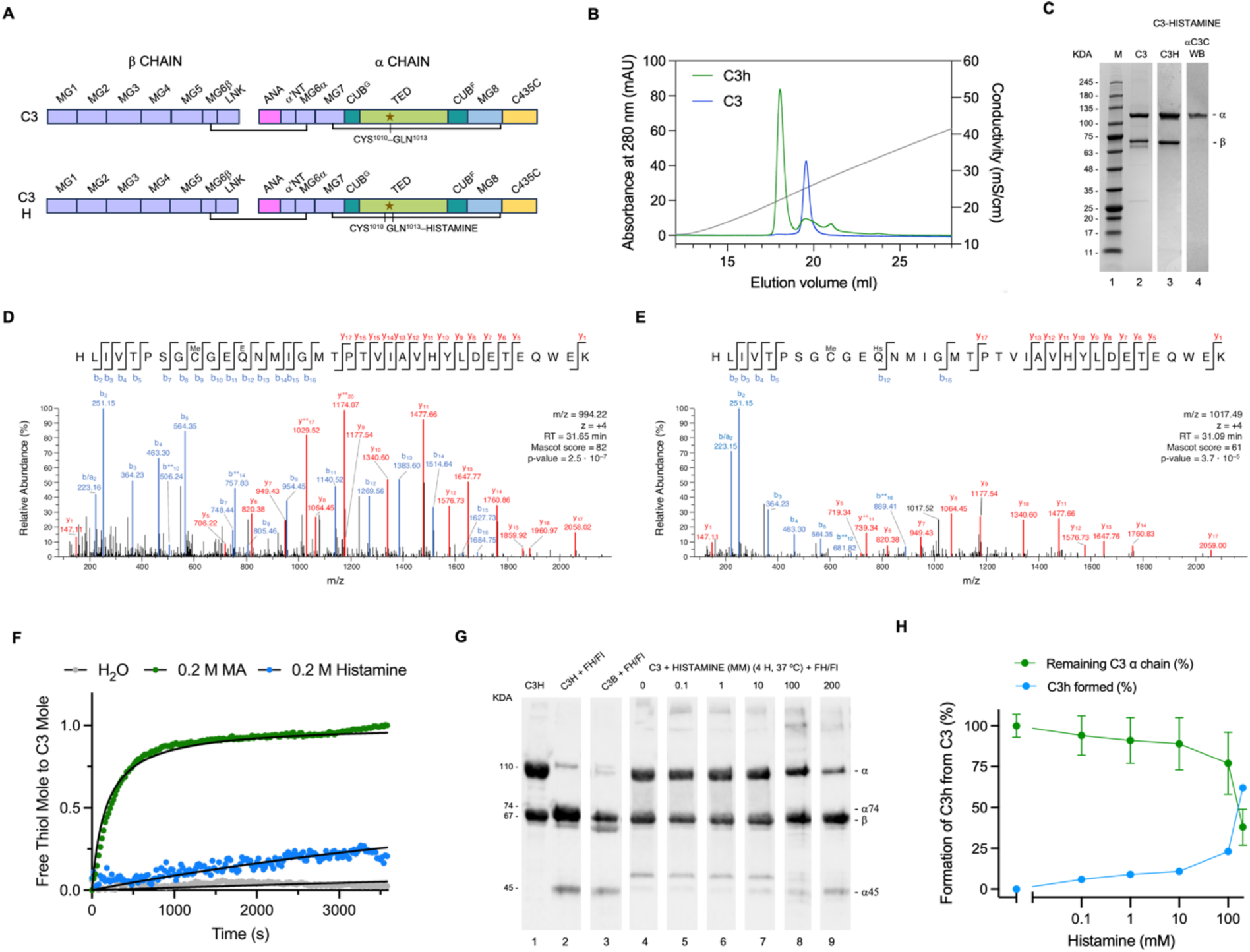
Histamine adduct of C3. (**A**) Schematic representation of the structural domains of C3 and C3h. The thioester bond residues Cys1010 and Gln1013 are marked with or without the histamine adduct. (**B**) Anion-exchange chromatography trace (280 nm) of control C3 (blue) and histamine-reacted C3 (green), well separated in a sodium chloride gradient (grey). (**C**) Coomassie-stained SDS-PAGE electrophoresis of control C3 (lane 2) and histamine-reacted C3 (C3h; lane 3), and Western blot of C3h with an anti-C3c mAb that detects the β chain of C3 (lane 4). (**D**,**E**) nLC-MS/MS identification of the thioester bond-containing peptide (1002–1036) in tandem MS/MS ion fragmentation spectra for control C3 (**D**) and after histamine modification of Gln1013 (**E**). The peptide sequence is shown, with the main b-series and y-series ion fragments labelled on the sequence and the spectra. The free thiol group of Cys1010 was methylated by treatment with iodoacetamide, as indicated by the Me label (**D**). The presence of the histamine adduct on Gln1013 is indicated with the Hs label (**E**). (**F**) Progress curves of the spontaneous formation of C3(H_2_O) (control, grey) and the C3 adducts with 0.2 M methylamine (MA) or 0.2 M histamine at 37 °C. (**G**,**H**) Histamine dose-dependent conversion of C3 to C3h upon incubation for 4 h at 37 °C and treatment with FH and FI. Coomassie-stained SDS-PAGE electrophoresis of dose samples (**G**). Quantification of the rate of conversion by following the appearance of the α74/α45 chains (green, *N* = 3) (its complement is shown in blue), diagnostic of the iC3b species generated from C3h and C3(H_2_O) (**H**).

To verify that the reaction had occurred at the thioester bond, we digested equal amounts of control C3 and C3h with trypsin and performed nano liquid chromatography-mass spectrometry on the sample digests (nLC-MS/MS). The MS/MS spectra identified 314 peptides from C3, representing a 94% sequence coverage, and 309 peptides from C3h, representing a 93% sequence coverage. We then focused our attention on the tryptic peptide comprising amino acids ^1002^HLIVTPSGCGEQ(deamidated)NMIGMTPTVIAVHYLDETEQWEK^1036^, which encompasses the thioester bond residues and includes a Glu residue instead of the thioester Gln1013 formed by the hydrolysis of the thioester bond. In C3, this peptide was detected with an experimental monoisotopic peptide mass of 3972.84 Da (theoretical monoisotopic mass of 3926.88 Da), and the excess +45.96 Da was ascribed to the methylation of the Cys1010 side chain sulfur atom. In contrast, in C3h, the equivalent peptide bearing a histamine moiety covalently bonded to Gln1010 side chain was detected with an experimental monoisotopic peptide mass of 4065.91 Da (theoretical monoisotopic mass of 4022.02 Da), and the excess +43.89 Da was again ascribed to the methylation of the Cys1010 side chain sulfur atom. The peptide mass shift between the two monoisotopic peptide masses is 93.07 Da, corresponding to the monoisotopic mass of histamine, less the amino terminal moiety (94.06 Da) (**Fig. 1D,E**).

These results show that C3 forms a covalent adduct with histamine through the net addition of histamine to the side chain of Gln1013 in the thioester bond.

### C3h forms at histamine pathophysiological concentrations *in vitro*

Next, we investigated the rate of formation of C3h from its C3 precursor in the presence of both saturating and pathophysiologically relevant concentrations of histamine. At the saturating concentration of 200 mM histamine, necessary to drive the complete histaminylation of C3 in a convenient time frame, 50% of C3 would be converted to C3h in 2.2 h (**Fig. 1F**). While the smaller primary amine methylamine reacts more quickly (half-life of ∼5 min under the same conditions), the reaction with water proceeds so slowly than it cannot be measured in the assay (**Fig. 1F**).

To evaluate the effect of pathophysiologically relevant concentrations of histamine, we incubated C3 at the physiological concentration of 1 mg/ml (∼5.4 µM) with various histamine concentrations from 200 mM to 1 µM for 4 h at 37 °C. Then we evaluated the amount of C3h generated from C3 by reaction with FH and FI. While C3 is not cleaved by FI, for C3h and C3(H_2_O), FI cleavage yields the two proteolytic products α74 and α45 chains; in contrast, FI cleavage of C3b yields the α67 chain instead of the α74 chain due to the lack of C3a. As demonstrated by SDS-PAGE electrophoresis and Western blotting, C3h can be produced from native C3 at all histamine concentrations tested in our experiments in a histamine-dependent manner (**Fig. 1G,H**). At the highest histamine concentrations (100-200 mM), C3 was fully or nearly fully converted to C3h during the incubation. However, at the lower, more physiologic concentrations of histamine (1-1000 µM), the amount of C3 converted to C3h was significantly smaller, requiring detection by Western blotting. Notably, after 4 h of incubation with 100 µM histamine, a pathophysiologically relevant concentration in tissues after mast-cell degranulation, approximately 6% conversion of C3 to C3h was achieved. This seemingly low conversation rate, however, should be high enough to cause a significant pathological deregulation of the AP complement self-amplification cycle (see **Fig. 2C**).

**Fig. 2.**
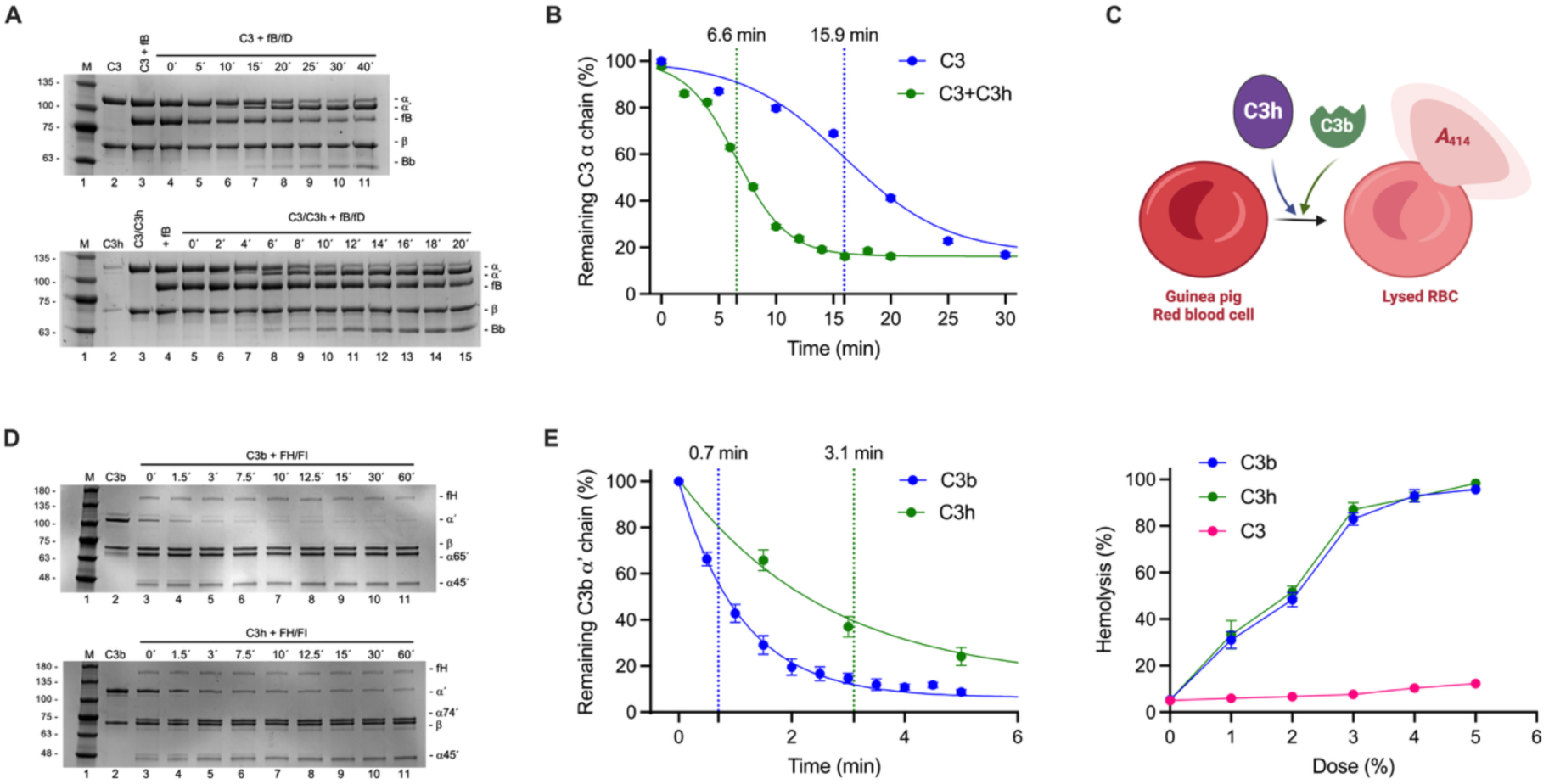
C3h forms an active C3 convertase downregulatable by FH and FI. (**A,B)** C3h accelerates C3 cleavage by FB and FD. Time course of C3 activation by FB and FD at 37 °C and time course of C3h-catalyzed C3 activation by FB and FD (1:10 molar ratio) under otherwise identical conditions (**A**), and (**B**) fitting of the intensity of the remaining C3 α chain in (**A**) to a 4-parameter log-logistic regression function. Half-lives of C3 are annotated and marked with vertical dotted lines. All experiments were performed at least in triplicate. (**C**) Progress of the hemolysis of guinea pig erythrocytes followed by the increase in absorbance at *A*_414_ following the release of the heme group. C3b was used as a positive control for lysis. Percent hemolysis calculated as the mean ± s.e.m. of three (*N* = 3) independent experiments. (**D**,**E**) FH-assisted FI cleavage of C3b into C3c and C3dg, and of C3h into C3a-containing C3c and histamine-containing C3dg (**D**), and (**E**) fitting of the intensity of the remaining C3b or C3h α’ chain in (**D**) to a 4-parameter log-logistic regression function. Half-lives of C3 are annotated and marked with vertical dotted lines. All experiments were performed at least in triplicate.

In conclusion, C3 can react with histamine *in vitro* to generate C3h at histamine concentrations previously measured in tissues under physiologic and pathologic conditions.

### C3h assembles a functional fluid-phase AP C3 convertase

We hypothesized that C3h might be able to assemble functional C3 convertases to initiate the self-amplifying loop of the alternative pathway in the fluid phase, as has been demonstrated previously for C3(H_2_O). To test this idea, we first assessed the ability of C3h to catalyze either C3 or its own cleavage in the presence of FB and FD (**Fig. 2**). Although C3h sustained C3 activation, C3h itself did not undergo cleavage with FB and FD, indicating that the conformation of C3h is incompatible with being a substrate for the AP C3 convertase (**Fig. 2**). Furthermore, in C3 activation assays with FB and FD, the presence of a small amount of C3h (1:10 molar ratio to C3) accelerated C3 cleavage by a factor of 2.4 (Welch’s t-test, *P* = 6 x 10^-8^), thereby indicating a strong catalytic effect (**Fig. 2A,b**). These results support the notion that C3h is equivalent to C3(H_2_O) or C3b in its capacity to assemble functional AP C3 convertases and its unsuitability to serve as a substrate for the same enzyme complex.

We then tested whether small amounts of C3h could start the self-amplification loop in fluid phase and promote complement activation on surfaces in a hemolytic assay (**Fig. 2C**). We used guinea pig erythrocytes as they are particularly valuable in complement hemolytic assays, because of their high sensitivity to complement, cross-species compatibility, and reliable and reproducible lysis. We calibrated the hemolytic assay so that the baseline lysis level is 20%. Purified C3b was used as a positive lysis control, as C3b(H_2_O) can efficiently activate complement factors in solution to induce reactive hemolysis effectively. We then titrated C3h in the same assay, finding that C3h behaves qualitatively and quantitatively as C3b(H_2_O) (**Fig. 2C**). These experiments indicate that small quantities of *in-situ* produced C3h have the potential to initiate the self-amplification loop of the AP and cause cell hemolysis in an *in vitro* system.

Finally, we investigated whether C3h can be inactivated by the FH cofactor activity of FI, a property shared by C3(H_2_O) and C3b that is essential to maintain a low yet significant level of basal complement activation. In standard assays at 37 °C with FH and FI, C3h was converted into C3a-containing C3c and C3dg at a rate that is 4.5 times slower than that of C3b (Welch’s t-test, *P* = 0.001), based on the time to reach 50% turnover, which was 0.7 min for C3b and 3.1 min for C3h (**Fig. 2D,E**). Therefore, in plasma and tissues, the half-life of C3h must be longer than that of C3(H_2_O), though still in the range of minutes. Furthermore, these results indicate that FH and FI are responsible for breaking down the C3h that may spontaneously arise in plasma or tissues, as they do with C3(H_2_O), resulting in a steady state characterized by a low-level activity necessary to prime the AP complement activation.

Taken together, these findings prove that C3h is a fully functional form of C3 that resembles C3(H_2_O) and C3b in its competence to assemble a catalytically active AP C3 convertase and cause C3 activation in fluid phase as well as reactive cell lysis, and in its susceptibility to be dampened by fluid-phase complement regulators.

### C3h is labile under physiological conditions and spontaneously releases C3a

As we observed that C3h could spontaneously cleave into C3hb (C3b-histamine adduct) and C3a much more readily than C3, we sought to investigate the stability of C3h under various conditions. First, we carried out a trypsin susceptibility assay whereby C3 or C3h is subjected to limited proteolysis with trypsin (1:100 (w/w)). In these assays, we observed that C3h could be cleaved into C3hb and C3a at a rate 9.5 times as fast as C3 with half-lives of 1.1 min versus 10.5 min, respectively (Welch’s t-test, *P* = 4 x 10^-5^) (**Fig. 3A,B**). These results suggest that the sequence of the α chain (residues 672–1663) connecting the anaphylatoxin domain (residues 672–747) with the α’ chain (residues 748–1663) has become sensitized to protease cleavage by a conformational change triggered by the reaction with histamine.

**Fig. 3.**
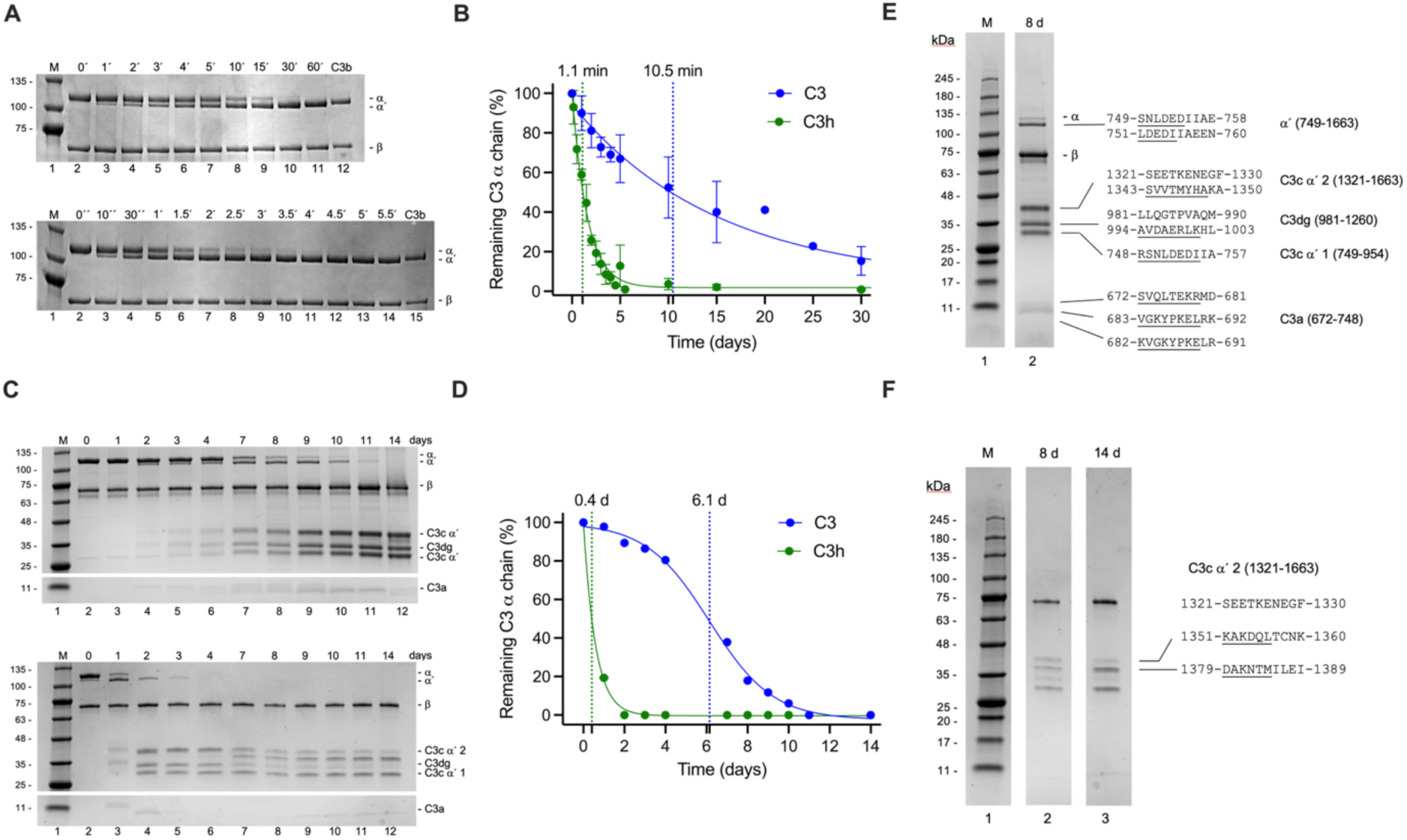
C3h is a labile species that spontaneously cleaves off into C3b-histamine and C3a. (**A**,**B**) Trypsin susceptibility assays of C3 (**A**, top) and C3h (**A**, bottom) with 1:100 (w/w) trypsin, analyzed by Coomassie-stained SDS-PAGE electrophoresis; (**B**) fitting of data in (**A**) to an exponential decay function. Half-lives are annotated and marked with vertical dotted lines. All experiments were performed at least in triplicate. (**C**,**D**) Spontaneous chemical cleavage of C3 (top, **C**) and C3h (bottom, **C**); (**D**) fitting of data in (**C**) to a 4-parameter log-logistic function (C3) or an exponential decay function (C3h). Half-lives are annotated and marked with vertical dotted lines. (**E,F)** N-terminal sequencing of spontaneous chemical cleavage products of C3 (**E**) and C3h (**F**).

Second, we incubated C3 and C3h for two weeks at 37 °C in a physiological phosphate buffer and sampled the reaction progress at time intervals. We added protease inhibitors to ensure that no residual protease activity could contaminate the results. Analysis of the spontaneous chemical cleavage products by SDS-PAGE gel electrophoresis and band quantification revealed that C3h was cleaved off far faster than C3, with half-lives of 0.4 d (9.6 h) versus 6.1 d, respectively, although the band degradation pattern was similar (**Fig. 3D,E**). In both cases, the main degradation products of the α chain, C3a and the α’ chain, were fast trimmed or broken down into other smaller fragments (**Fig. 3C,E,F**). At least 2-3 distinct bands corresponding to C3a could be observed during the spontaneous cleavage of C3 and C3h (**Fig. 3C**). The other main degradation product of the α chain, the α’ chain, rapidly broke down into the well-known fragments C3c α’ 1 (residues 749–954), C3d(g) (C3dg encompasses residues 955–1303, of which residues 955–1001 correspond to C3g and residues 1002–1303 to C3d), and C3c α’ 2 (residues 1321–1663) (**Fig. 3C,E,F**).

Interestingly, the main time-independent difference between the spontaneous degradation pattern of C3 and C3h concerned C3d(g) and C3c α’ 2 (**Fig. 3C**), which represent the domain harboring the histamine adduct and the region immediately following it, respectively. While when derived from C3 both protein fragments remain essentially stable as judged by SDS-PAGE electrophoresis for up to 14 d, the corresponding fragments derived from C3h underwent additional chemical degradation (**Fig. 3C**). Within 2 d, the C3h-derived α chain converted into the α’ chain by losing C3a, and the former fully degraded into C3c α’ 1, C3d(g), and C3c α’ 2. From day 7 onward, C3c α’ 2 lost a ∼30-amino-acid fragment and stabilized into a 37 kDa protein fragment (**Fig. 3C**). Submitting these degradation products to N-terminal sequencing confirmed the virtually authentic nature of, as well as the differences between, the spontaneous chemical cleavages (**Fig. 3E,F**). Thus, the α’ chain derived from C3 or C3h started with either the authentic sequence or lost at most two amino acids (**Fig. 3E,F**). The stable N terminus of C3-derived C3d(g) started at 994, closer to the canonical start of C3d (1002) than to that of C3dg (955) (**Fig. 3E**). In stark contrast, C3h-derived C3d(g) quickly degraded after reaching its peak concentration at 2 d (**Fig. 3F**). As for C3h-derived C3c α’ 2, the N-terminal sequences of the earlier and later products started at 1351 or 1379, respectively (**Fig. 3F**). The longest of the two C3c α’ 2 fragments derived from C3h was still 12-amino-acids shorter than that derived from C3 (**Fig. 3E,F**), further suggesting that the C3h-derived fragments were more labile.

In conclusion, C3h appears to be significantly more labile than C3 and readily undergoes spontaneous chemical cleavage at 37 °C, releasing both C3hb and the C3a anaphylatoxin.

### Structure of C3h

To provide direct evidence for the histamine adduct and interrogate the structural changes undergone by C3 upon reaction with histamine, we determined the X-ray crystallographic structure of C3h to 2.81-Å resolution (**Fig. 4**) from crystals obtained from concentrated C3h (5 mg/ml). The diffraction data were phased by molecular replacement using two structures of C3b (PDB 2I07 and 5FO7) as the source of the search models and considering the placement and orientation of the C345c and thioester-containing domain (TED) domains independently. Cycles of macromolecular refinement and manual building were performed until convergence. The final refined model displayed excellent refinement statistics and correct stereochemistry (**Table 1**).

**Fig. 4.**
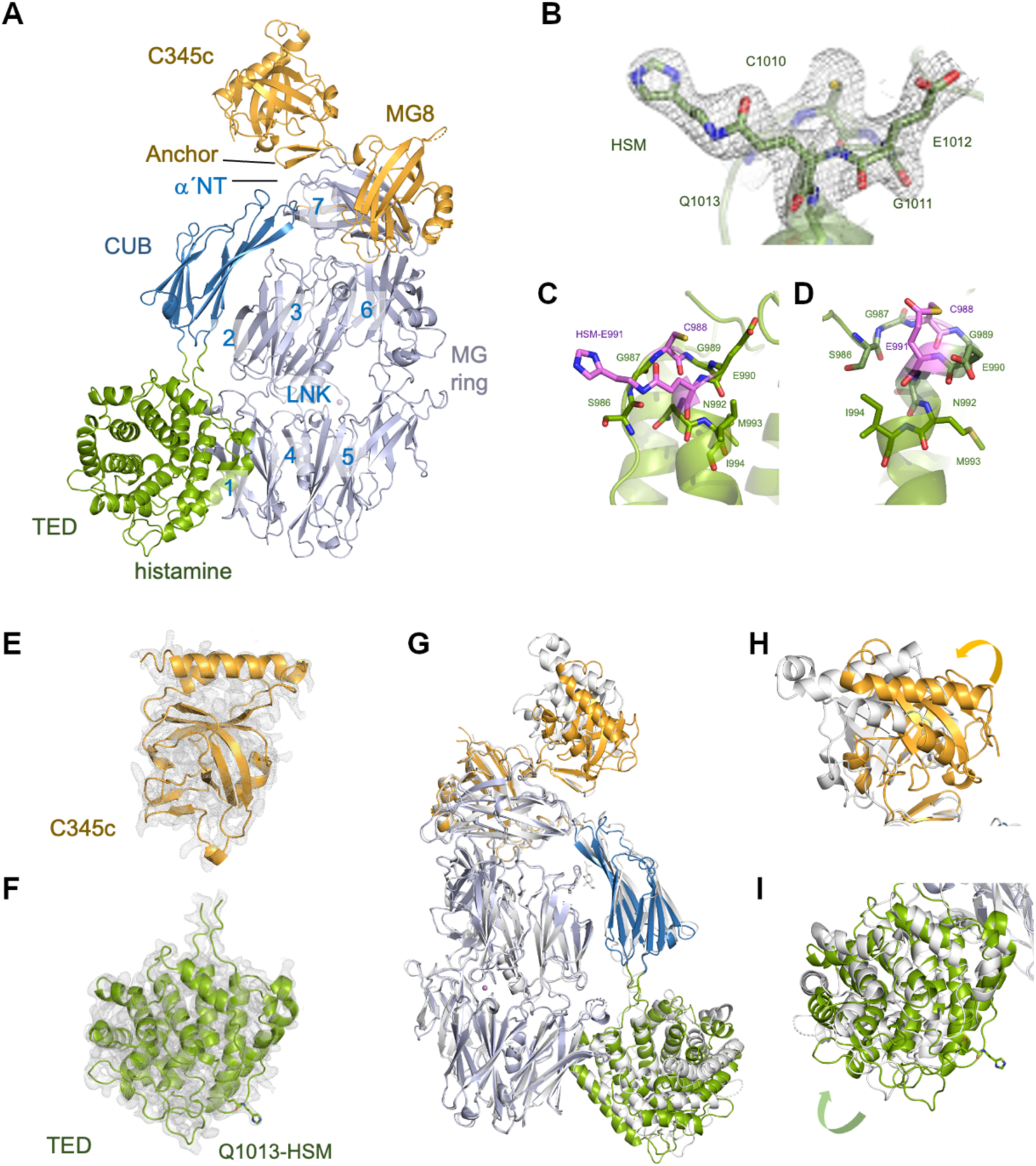
3D structure of C3h. (**A**) Cartoon representation of C3h. Canonical domains, regions, and motifs in C3 are annotated and color-coded. (**B**) Zoom into the histamine adduct and thioester bond residues Cys1010–Gln1013. The final refined σ_A_-weighted 2mFo-DFc map is shown. (**C**,**D**) Conformation adopted by the thioester bond residues after the histamine adduct formation (C) and in C3 before the thioester bond breakdown (**D**). (**E**,**F**) Quality of the σ_A_-weighted 2mFo-DFc map for the domains with the most changes or flexibility: C345c (**E**) and the TED domain (**F**).

**Table 1.** X-ray crystallographic statistics.

|  |  |
| --- | --- |
|  | **C3h** |
| Synchrotron beamline | BL13-XALOC (ALBA) |
| **Data processing statistics** |  |
| Wavelength (Å) | 0.97926 |
| Resolution range (Å) | 45.61-2.81 (2.86-2.81)* |
| Space group | *P*1 |
| Unit cell dimensions |  |
| *a*, *b*, *c* (Å) | 76.22, 91.98, 93.51 |
| $\alpha$, $\beta$, $\gamma$ (°) | 113.83, 104.22, 100.96 |
| Total reflections | 180,099 (9,208) |
| Unique reflections | 50,574 (2,627) |
| Multiplicity | 3.6 (3.5) |
| Completeness (%) | 97.30 (95.48) |
| $\langle I \rangle / \sigma(\langle I \rangle)$ | 6.33 (0.69) |
| Wilson *B*-factor (Å<sup>2</sup>) | 78.41 |
| *R*<sub>merge</sub> | 0.1306 (1.623) |
| *R*<sub>meas</sub> | 0.1543 (1.913) |
| *R*<sub>pim</sub> | 0.0812 (1.000) |
| CC<sub>1/2</sub> | 0.993 (0.401) |
| CC* | 0.998 (0.756) |
| **Refinement statistics** |  |
| Reflections used | 50,446 (2,746) |
| Reflections used for *R*<sub>free</sub> | 2,520 (135) |
| *R*<sub>work</sub> | 0.2036 (0.3747) |
| *R*<sub>free</sub> | 0.2464 (0.4197) |
| Non-H atoms | 12,364 |
| Macromolecules | 12,248 |
| Ligands | 70 |
| Solvent | 46 |
| Protein residues | 1548 |
| RMS(bonds) (Å) | 0.002 |
| RMS(angles) (°) | 0.55 |
| Ramachandran favored (%) | 99.42 |
| Ramachandran allowed (%) | 0.58 |
| Ramachandran outliers (%) | 0.00 |
| Rotamer outliers (%) | 0.07 |
| Clashscore | 5.04 |
| Average *B*-factor | 86.68 |
| Macromolecules | 86.52 |
| Ligands | 124.24 |
| Solvent | 72.12 |
| PDB code | 9SKN |

As mature C3, C3h consists of two chains stabilized by disulfide bridges, the β chain (residues 23–667) and the α chain (residues 672–1663). The structural organization of the 13 domains in C3h can be described as a central β-ring body in the shape of a keyholder that includes the macroglobulin (MG) 1-6 domains and the linker (LNK) domain, the shoulders (domains MG7 and MG8), a neck (the Anchor region), a head (C345c domain), and an arm (“complement C1r/C1s, Uegf, Bmp1” (CUB) domain) holding the globular TED domain. C3b is generated from C3 by the proteolytic excision of the small anaphylatoxin domain (residues 672–748) by the serine proteases FB and FD. In contrast, in C3h, the domain rearrangements following histamine reaction with the C3 thioester moiety initially preserve the integrity of the proteolytic cleavage site connecting C3a. However, in our crystal structure, C3h has lost the C3a anaphylatoxin domain by spontaneous chemical cleavage (**Fig. 4A**). This cleavage results in a shortened form of the α chain termed α’ chain, akin to that of C3b (residues 749–1663), and the spontaneous release of the entire C3a (672–748) (**Fig. 3C**). The structural analysis of C3h revealed a conformation most closely resembling that of the C3 activated product C3b, with a root mean square distance (RMSD) of 2.8 Å (1544 Cα atom pairs) after superposition (**Fig. 4A,G-I**). Restricting the superposition to the TED domain improved the RMSD after superposition to 0.66 Å (all 276 Cα atom pairs), indicating that the TED domain of C3h has adopted a fully activated conformation(*24*) (**Fig. 4H**).

Before refinement was started, omit maps centered around the residues forming the thioester moiety of C3 revealed clear electron density for the histamine adduct covalently bound to Gln1013 (**Fig. 4B**). Inspection of the structural changes in the TED domain of C3h around the Gln1013-histamine covalent adduct revealed the conformational adjustments present in the fully activated C3b(*24*) (**Fig. 4F,G,I**). The zig-zag arrangement of helices α0 (966-974) and α1 (977-982) and the kinked backbone conformation of helix α2 (990-1009) characteristic of the TED domain of C3 have unraveled into an extended conformation in C3h, where α0 disappears into a more elongated conformation and α1 and α2 straighten toward a more relaxed structure (**Fig. 4A,F,G**). These changes are paralleled by changes in the loop spanning residues 1268–1278 between helices α15-α16, which rest on helices α0-α1 in C3. At the same time, in C3h, helices α2-α5 move inwardly by a maximum of 9.6 Å as they rotate by ∼16° against the N-terminal segment of the TED domain. In turn, these changes correlate with an increase in the distance between the thioester bond atoms from 2.8 to 6.4 Å, an extension that is facilitated by the flexible Gly residues nearby the thioester residues (Gly1009 and Gly1011). Once the Gln1013-histamine adduct forms and the conformational changes necessary for activation are complete, the histamine side chain becomes fully exposed to the solvent, sticking ∼5.5 Å above the molecular surface of the TED domain and apparently lacking specific interactions with the rest of C3h (**Fig. 4A-C**).

### Reaction mechanism of nucleophilic attack on C3 thioester

To shed light on the chemical reaction mechanism of the tick-over reaction with water or histamine as the nucleophiles, we performed exhaustive DFT/MM free energy simulations of both mechanisms. To model the reaction in the C3 structure we aligned the TED domain of C3h, including the histamine covalent adduct, with that of C3 (PDB 2A73), resulting in the structure presented in **Fig. 5A**. After equilibration (**Supplementary Fig. S4**), we used the resulting structure to trace the Minimum Free Energy Path (MFEP) for the reaction between the thioester bond and histamine, from products to reactants. We then changed histamine by a water molecule to study the hydrolysis of the thioester bond. The free energy profiles of the two reactions are presented in **Fig. 5B**, while **Fig. 5C-H** display the structures of reactants, transition states (TSs), and products for both processes.

**Fig. 5.**
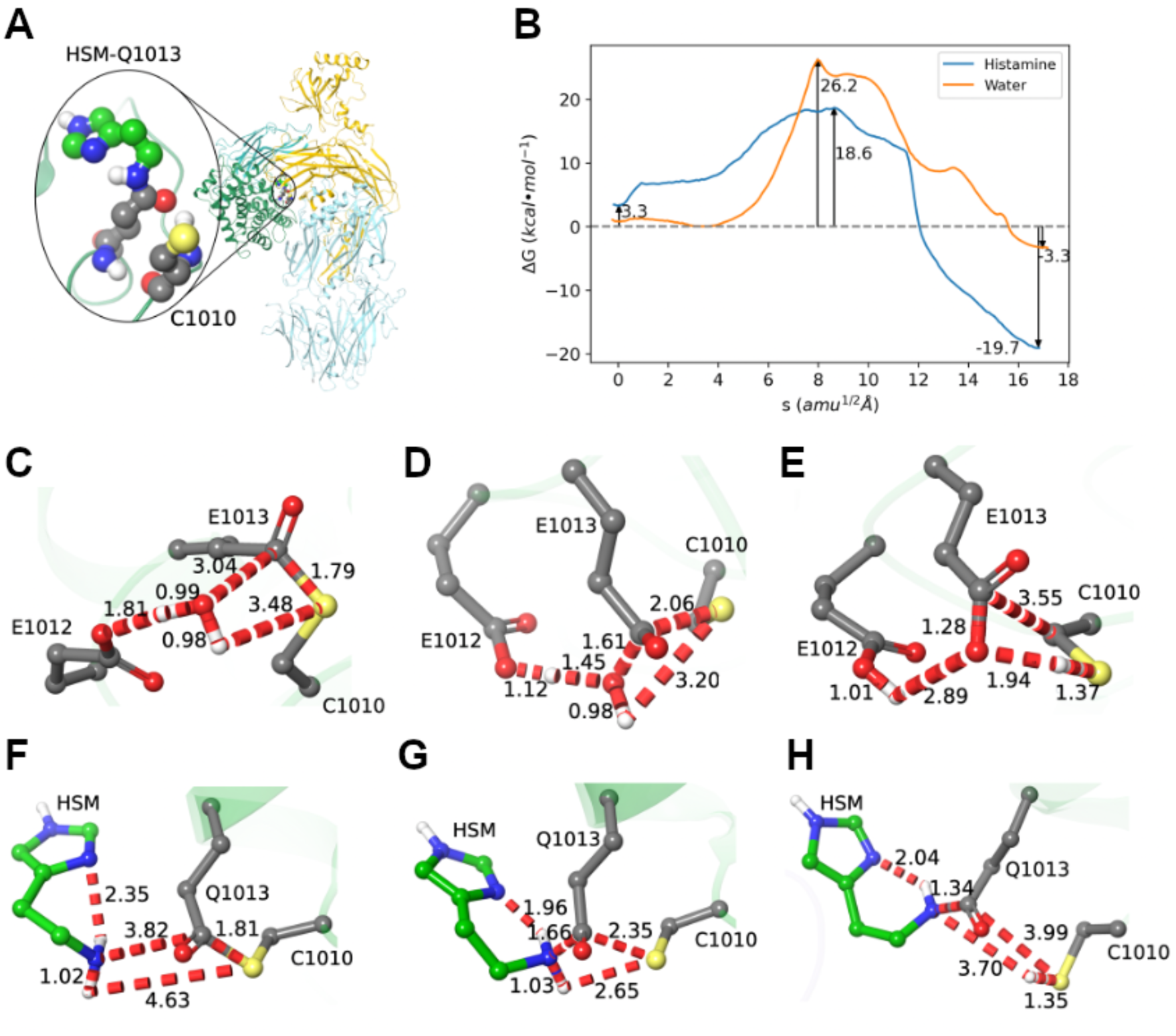
The tick-over reaction mechanism with water and histamine as the main nucleophiles. (**A**) Adduct structure prepared in a C3-like protein conformation as a model of the tick-over reaction products. (**B**) Free energy profiles corresponding to the hydrolysis and aminolysis reactions. Activation and reaction free energies are displayed in the graphic. (**C-E**) Structures corresponding to the reactants (**C**), transition (**D**) and product states (**E**) for the reaction of the thioester adduct with water. Carbon, sulfur, and oxygen atoms are displayed in grey, yellow, and red, respectively. Only water hydrogen atoms are shown (in white) for the sake of clarity. (**F-H**) Structures corresponding to the reactants (**F**), transition (**G**), and product states (**H**) for the reaction of the thioester adduct with histamine. Protein carbon, sulfur, and oxygen atoms are displayed in grey, yellow, and red, respectively, while histamine atoms are shown in green (carbon) and blue (nitrogen). Only polar histamine hydrogen atoms are displayed (in white) for the sake of clarity.

The cleavage of the thioester bond by a water molecule is assisted by a nearby glutamate (Glu1012). This residue orientates the water molecule for the nucleophilic attack (**Fig. 5C**) and activates it abstracting one of its protons (see TS structure in **Fig. 5D**). After breaking the thioester bond the sulfur atom of Cys1013 is protonated leading to the final products (**Fig. 5E**). The evolution of the bond breaking/forming distances during the reaction path is detailed in **Supplementary Fig. S5** and **S6**. The free energy profile (**Fig. 5B**) indicates that this is a spontaneous process (with a reaction free energy of -3.3 kcal·mol^-1^), but with slow kinetics. The activation free energy is 26.2 kcal/mol, which, according to Transition State Theory, can be translated into a reaction half-life (*t*_1/2_) at 300 *K* of about 15 d. This theoretical estimation agrees with the experimental observation that hydrolysis of the thioester bond in C3 is very slow(*25*) and also with the fact that C3 is cleaved off with a half-time larger than 6 days (see above).

When we shifted our attention to histamine as the primary nucleophile, it became apparent that two opposing factors contributed most significantly to the mechanism. First, histamine at pH 7.4 is predominantly found in the monocationic form, with the terminal amino group protonated (the p*K*_a_ being 9.76)(*26*), which is unable to carry out the nucleophilic attack. Using an alchemical transformation (see Methods), we evaluated the free energy cost of deprotonating the terminal amino group to form the free base in the active site of C3 to be 3.3 kcal/mol (**Supplementary Fig. S7**). However, the stronger nucleophilicity of the neutral histamine amine compensated for this energetic penalty, leading to a thioester bond cleavage that, according to the free energy profile (**Fig. 5B**), is much more exergonic (with a reaction free energy of –19.7 kcal·mol^-1^) and several orders of magnitude faster than with water. The total activation free energy, including the cost of forming neutral histamine, is only 18.6 kcal·mol^-1^, which translates into a *t*_1/2_ for this chemical step of ∼4 s, about 3·10^5^ times faster than with water. The reason for this significantly enhanced activity is not only the more basic character of the amino group with respect to water, but also the formation of an intramolecular hydrogen bond with the Nδ atom of the histamine’s imidazole ring that enhances the nucleophilicity of the amine, as seen at the reactants structure in **Fig. 5F**. At the TS (**Fig. 5G**), the thioester bond cleavage is significantly more advanced than in the reaction with water (**Fig. 5D**). When the product is formed (**Fig. 5H**) the leaving cysteine is protonated, and the new C-N bond is fully formed. The complete evolution of the bond-breaking and forming distances during the aminolysis reaction path is detailed in **Supplementary Fig. S6**. Therefore, based on our DFT/MM calculations, while the chemical step would be rate-limiting when C3 is cleaved by water to produce C3(H_2_O), this would not be the case for C3h formation with histamine. In the conversion from C3 to C3h, the slowest step could be the conformational rearrangement of the TED domain.

### Dynamics of C3h and C3(H_2_O) support the lysis of the linker region

To provide additional structural information from intact C3h in solution, we resorted to SEC-SAXS, a technique that allows us to infer the size and hydrodynamic parameters of proteins in solution as the protein samples elute off a connected SEC column, and we used C3h freshly prepared to reduce the chance of spontaneously losing C3a (**Fig. 4D**). 3D shape restoration by SAXS *ab initio* methods allowed us to build a model of C3h containing C3a by rigid-body fitting of the C3a-less C3h crystallographic structure and then the structure of C3a (PDB 2A73) into the SAXS shape (**Fig. 4E**). To obtain an atomistic model of C3h based on the SAXS data, we performed rigid-body refinement of the model considering a fixed C3bh moiety and a movable C3a, resulting in a model with an excellent agreement to the experimental data (χ^2^ = 1.2) (**Fig. 5C,D**).

Using the SAXS-guided rigid body model of a C3a-containing C3h, we generated a complete model of C3h with MODELLER (**Fig. 6A**). We subjected it to a 1.1-µs molecular dynamics (MD) simulation to probe the statistical relevance of SAXS-compatible conformations and the conformational dynamics of C3a. During the trajectory, C3a adopted a series of closely related orientations that retained significant similarity with the SAXS data (**Fig. 6B**). Accordingly, the mean of the goodness-of-fit χ^2^ statistic for all the frames in the trajectory was 1.5 ± 0.2 (range = 1.06–2.25). The 25 conformations with the lowest χ^2^ values (range = 1.06–1.13) were extracted and inspected. All of them were closely related, showing C3a lying perpendicular to the C345c domain of C3h. In contact with it and the upper surfaces of the Anchor and MG8 domains (**Fig. 6C**). The residues connecting the C terminus of C3a with the cleavage site and following residues in the α chain (residues 743–753 of the α’NT segment) are highly solvent exposed, as shown by the decrease in buried surface area in this region between the averaged 25 lowest χ^2^ value conformations (573.4 Å^2^) and the reference structure of C3 (PDB 2A73) (1121.6 Å^2^) (Welch’s t-test, *P* < 10^-6^), or a 51% reduction in buried surface area (**Fig. 6B**). Likewise, the MD trajectory frames contained C3h structures with increased solvent exposure of the linker between C3b and C3a (Welch’s t-test, *P* < 10^-4^), regardless of how closely they reproduce the SAXS scattering (Welch’s t-test, *P* = 0.28). As anticipated, C3a is more solvent-exposed in both the C3a-containing C3h model and the MD trajectory compared to the C3 crystal structure, due to the significant conformational changes that accompany C3 activation (Welch’s t-test, *P* = 0.02) (**Fig. 6B**).

**Fig. 6.**
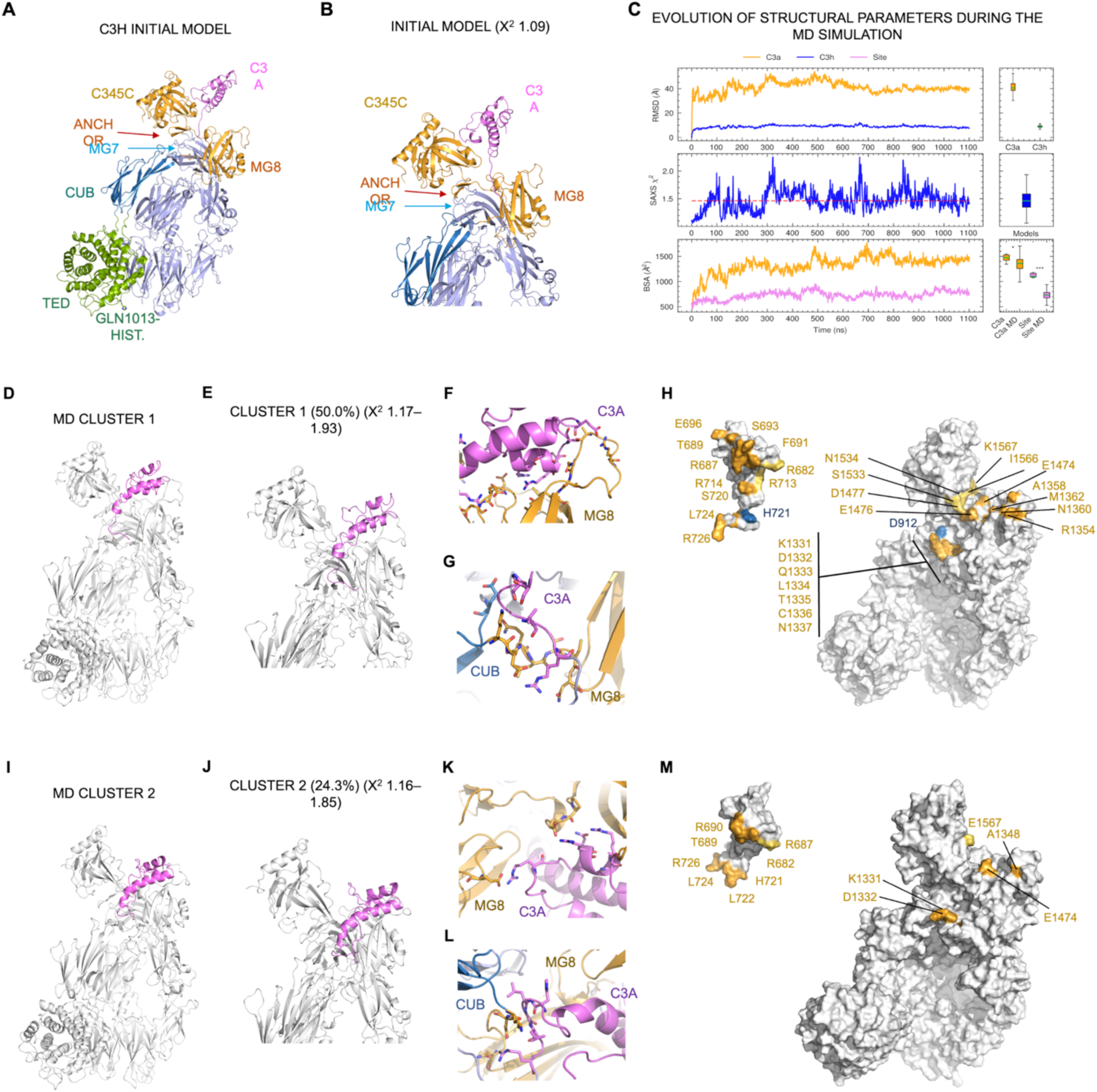
Dynamics of C3h/C3(H_2_O). (**A**,**B**) Complete model of C3h including C3a (in red) that represents the starting conformation for the 1.1-µs MD simulation (**A**) and zoom showing the starting position and orientation of C3a (**B**). (**c**) Progress of diagnostic variables during the MD simulation of C3h, including C3a. RMSD, root mean square deviation (Å) of C3h and C3a. SAXS goodness-of-fit χ^2^, with the mean value (1.46) marked by a dashed red line. BSA, buried surface area (Å^2^) calculated for C3a and for the residue segment targeted by proteolytic or spontaneous cleavage (Site: residues 742-752). Boxplots summarize the distribution of the variables of interest. The median, interquartile rank, and the range are shown. In the BSA boxplot, a comparison between the BSA values obtained for a sequence-complete model constructed from the C3 crystal structure (PDB 2A73) and the full simulation. Statistical significance of the mean difference is shown graphically (Welch’s t-test; *, *P* < 0.05; **, *P* < 0.01; ***, *P* < 0.001; ns, not significant). (**D-H**) Structural model of the most populated MD conformer is cluster 1 (50%). Overall structure (**D**) and zoom (**E**) of cluster 1 in cartoon, with C3a in violet. Details of the two main interfaces where there are contacts between C3a and the rest of C3 (**F**, **G**). Mapping of interacting residues in (**F**, **G**) on the molecular surface of either C3a or the rest of C3 is shown and annotated in (**H**). (**I**-**M**) As in **D**, **E**, **F**, **G**, **H**, respectively, for the second most representative MD conformer or cluster 2 (25%).

Next, we performed a structural similarity clustering algorithm on the MD trajectory to identify and classify the most frequently sampled spatial conformations. Using a 3 Å cutoff for RMSD calculations, the algorithm identified 17 distinct conformations, even though only the first 2–3 clusters were significantly populated (> 5%). The representative central structures of the initial model and the first two clusters (accounting for 50.0% and 24.3% of the whole MD trajectory) are shown in **Fig. 6c**. These highly abundant conformations indicate that C3a can stably reside wedged between the MG8, Anchor, and C345c domains upon C3 activation by hydrolysis or reaction with histamine. In this location, C3a is maximally removed from the area around the CUB, MG7, and MG6 domains, which undergo dramatic changes as C3 adopts a C3b-like conformation.

All combined, these structural data support a structure for C3h that fully exposes the region connecting C3h α chain and C3a to solvent nucleophiles or proteases, and a C3b-like structure for C3h after the loss of the C3a domain.

### Comparison of C3h with C3b in C3 (pro-)convertase

To provide insights into the capacity of C3h to assemble fully functional C3 convertases, we sought to compare the structure of C3h with that of C3b, as shown in the few structures of C3 pro-convertase and convertases available at present. Those structures include the C3 pro-convertase and its complex with FD (PDB 2XWJ, 2XWB), the complex of a dimeric form of the C3 convertase (C3bBb) with the small α-helical *Staphylococcus aureus* complement inhibitor (SCIN) (PDB 2WIN) and with the C3 convertase activator properdin. Superposition of these structures with C3h or C3a-containing C3h sheds insight into the structural basis for C3h and C3(H_2_O) assembly of fully functional C3 convertases (**Fig. 7**).

**Fig. 7.**
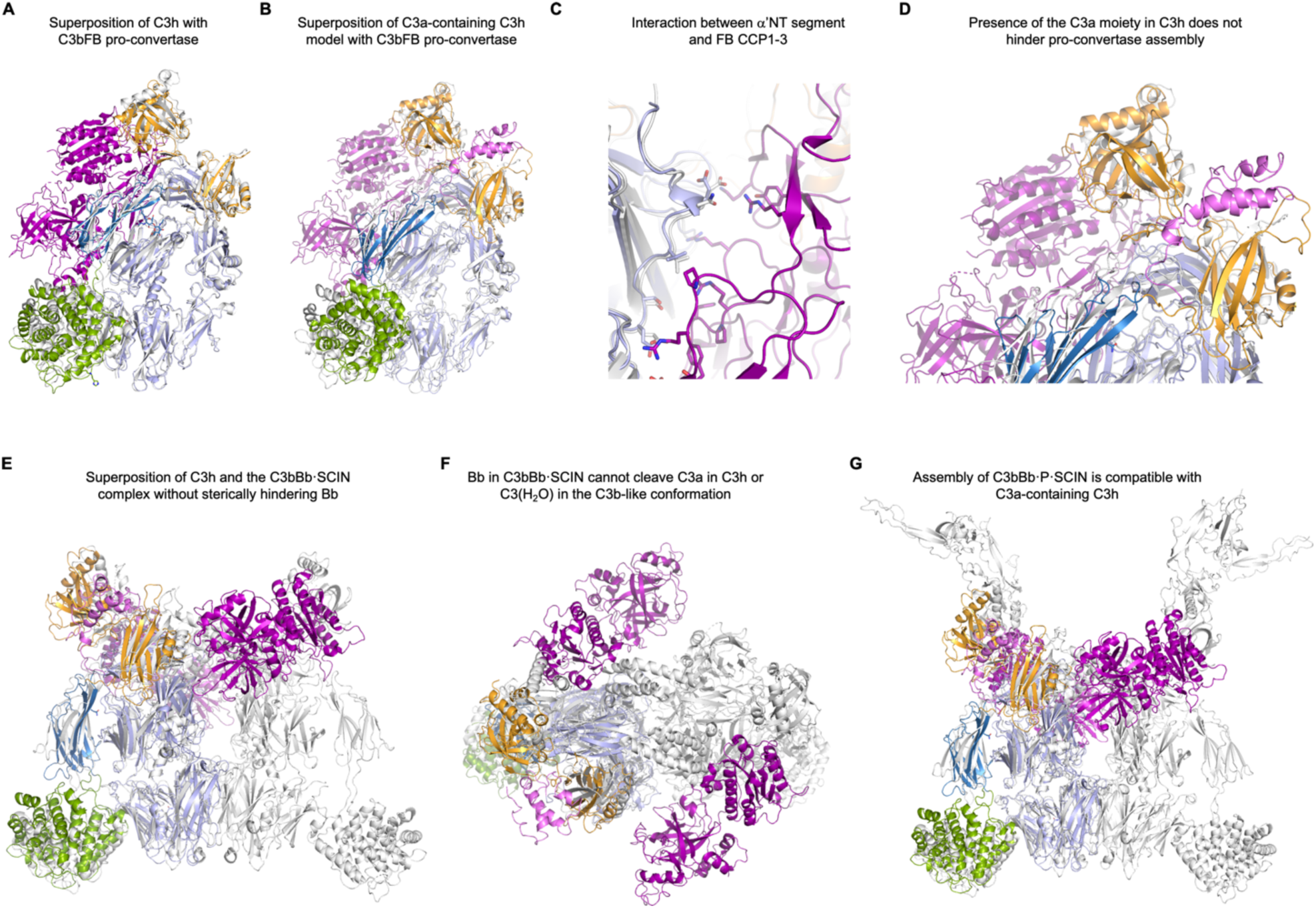
Structural implications of C3h/C3(H_2_O) for C3 pro-convertase and C3 convertase. (**A**) Superposition of C3h with the C3bB pro-convertase (PDB 2XWJ). C3h is shown in domain colors, all other structural components are shown in white (C3b) or pink (FB/Bb). (**B**) As in (**A**) with C3a-containing C3h. (**C**) Zoom into the α’NT segment of C3b/C3h in (**A**), showing that the segment makes few interactions with the CCP 1-3 domains of FB in the pro-convertase. (**C**) Zoom into the C345c domain of the C3bFB pro-convertase complexes to show that C3a in C3h cannot directly affect the complex assembly. (**D**,**E**) Side (**D**) and front (**E**) views of the structural comparison of C3h with the C3bBb·SCIN complex (PDB 2WIN). Here, the location of C3a on the opposite face to FB/Bb binding facilitates C3 convertase assembly. SCIN is shown in black and Bb in pink. (**F**) Superposition of C3a-containing C3h with the C3bBb·Properdin (P)·SCIN complex (PDB 6RUR).

Superposing C3h or C3a-containing C3h with the C3bFB pro-convertase structure (PDB 2XWJ) shows that C3a-less C3h can substitute for C3b in assembling a viable C3 pro-convertase by virtue of their highly similar architectures (**Fig. 7A,B**). The resultant C3 pro-convertase complex, which can be efficiently cleaved by FD (**Fig. 2D,E**), offers no steric impediment for the engagement of FD, the serine protease responsible for FB cleavage and activation to Bb that is the hallmark of the C3 convertase. Magnification of the superposed structures around the α’NT segment confirms that the surfaces involved in FB interaction are indistinguishable in the mature C3b/C3h structures (**Fig. 7C**). C3a-containing C3h, where the α’NT segment is still threaded inside the MG ring by the steric impediment provided by the presence of C3a, can still assemble a fully functional C3 (pro-)convertase, as the lack of the mature positioning of the α’NT segment contributes little to the stability of the complexes. Both the α’ and β chains of C3b/C3h make significant contributions to the extensive interaction with FB, but residues in the α’NT segment account for < 6% of all hydrogen bond interactions and ∼7% of the total buried surface area (672.8 Å^2^). In the structure of C3a-containing C3h or C3(H_2_O), where the α’NT segment remains threaded through the MG ring by being linked with C3a, the conformers observed during the MD simulations are confined to the opposite face of the C345c domain, in contact with the MG8 domain and the Anchor region (**Fig. 7D**). In these locations, C3a provides no steric impediment to either C3 pro-convertase assembly or the association with FD that precedes C3 convertase formation.

The available structures of the C3 convertase consist of dimeric forms stabilized by SCIN (PDB 2WIN) and further by properdin (PDB 6RUR). Although they are meant to approximate the active configuration of the enzyme complex, the presence of SCIN, a potent natural inhibitor, and the conformation of key active site residues in the active site of the SP domain of Bb indicate that the observed conformation is not consistent with a fully active C3 convertase. With this caveat in mind, our superposition of C3h or C3a-containing C3h with dimeric C3 convertase structures reveals that their assembly would be easily accomplished in the presence of C3a (**Fig. 7E-G**).

Hence, after the spontaneous or catalyzed loss of C3a, C3 convertases could be assembled from C3a-less C3h or C3(H_2_O) that are virtually indistinguishable from the authentic surface-attached C3 convertases.

### Structural implications of C3h for FH cofactor activity

By acting as a cofactor for the plasma serine protease FI, FH is the major negative regulator of the alternative pathway activation in fluid phase. Previous studies have shown that the CCP 1–4 domains of FH participate in C3 convertase inactivation by first associating with C3b across its entire surface, from the C345c domain to the bottom of the MG ring and the TED domain, and then bringing in the constitutively active FI that makes two consecutive cleavages in C3b. The first cleavage converts C3b into iC3b, a proteolytic fragment that loses the capacity of assembling C3 convertases, while the second cleavage splits iC3b into free C3c and surface-attached C3d(g).

The structure of C3h is compatible with the fluid-phase inactivation of C3h-containing C3 convertases in plasma by FH-mediated FI (**Fig. 8**). In the crystallographic structure of C3bFH (PDB 2WII) (*27*), the α’NT segment is sandwiched between the MG6–MG7 domains of C3b and the CCP 1–2 domains of FH. Although significant, the number of electrostatic interactions (four hydrogen bonds and one salt bridge) and the amount of buried surface area (403.7 Å^2^) contributed by the α’NT segment to the interaction with FH CCP 1–4 is far smaller than the rest of C3b (11 hydrogen bonds, 7 salt bridges, and 1619.8 Å^2^ buried surface area). As C3h is characterized by a more C3-like arrangement of the α’NT segment-anaphylatoxin domain, it lacks the complete interaction surface of C3b with FH CCP 1–4. In addition to possessing a complete docking surface for FH CCP 1–4, the C345c domain of C3b is relocated and reoriented upon FH interaction (**Fig. 8**). As the C3h structure is most similar to that of free C3b than to C3b in complex with FH CCP 1–4, the position and orientation of its C345c domain differs significantly from that of C3bFH (**Fig. 8**). However, the superposition with the C3a-containing C3h model reveals that, as with the C3 pro-convertase and C3 convertase complexes, C3a presents no steric impediment for the C345c domain and, therefore, its presence is not likely to prevent the required conformational change (**Fig. 8**).

**Fig. 8.**
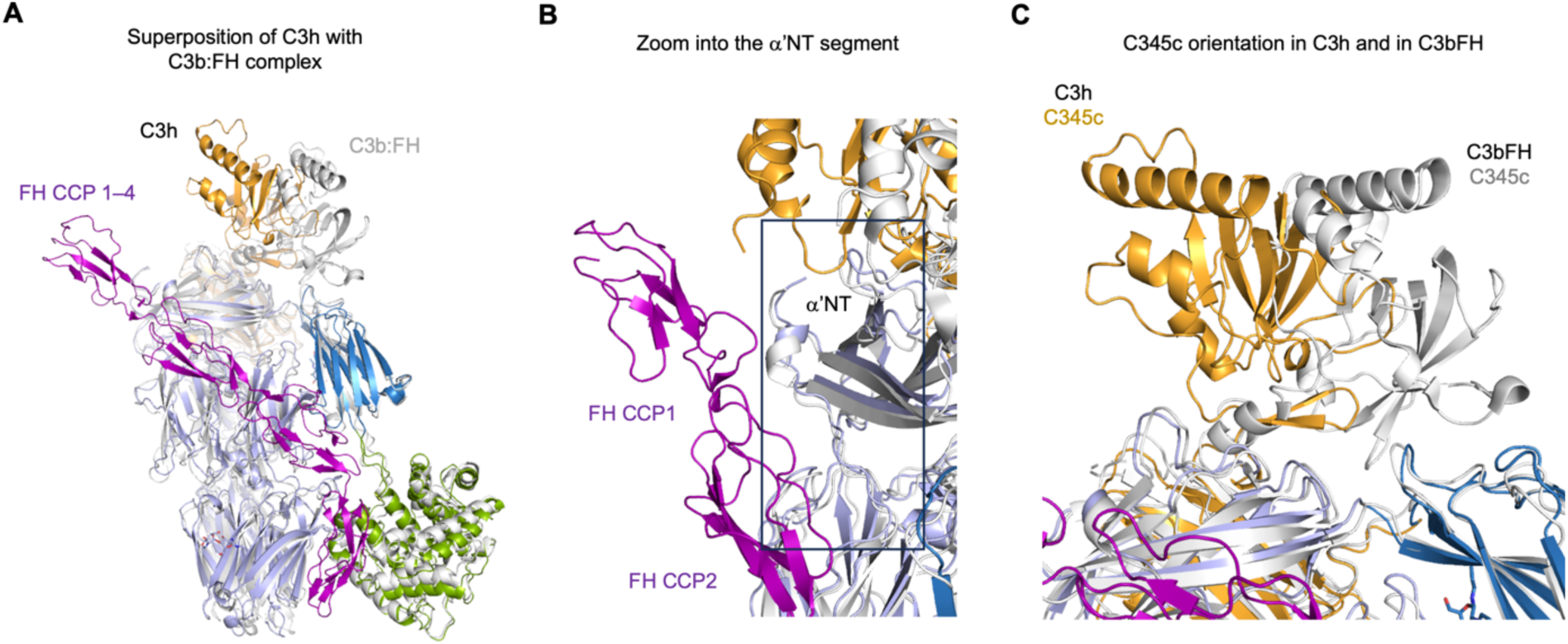
Structural basis of C3h convertase negative regulation by FH. (**A**) Superposition of C3h and C3bFH (PDB 2WII). C3h is shown in cartoon representation and domain colors. C3b:FH is in white (C3b) and purple (FH CCP 1–4). (**B**) Zoom into the α’NT segment of C3h/C3b, with the bounding box around it shown in black. (**C**) Zoom into the C345c domains of C3h/C3b to highlight the difference in position and orientation between free C3b/C3h versus C3b in complex with FH CCP 1–4.

All together, these results explain how FH/FI can rapidly inactivate C3h (and, consequently, also C3(H_2_O)) shortly after they appear in plasma. Even though this fast inactivation process cannot hinder the low level of basal activation necessary for the normal functioning of the alternative pathway, it ensures that under physiological conditions, activation does not spiral out of control.

### Quantitative modeling of histamine conjugation to C3

To develop a quantitative understanding of the concentrations that histamine-conjugated C3 species can reach under various pathophysiological situations, we used the C-model software, a comprehensive, enhanced PK/PD software specifically tailored to the complement system(*28*). To this end, we implemented in C-model equations to model histamine metabolism (synthesis and diamine oxidase (DAO)-mediated degradation), the histamine conjugation reaction with thioester-containing proteins (only for C3), as well as the subsequent activation and degradation reactions of C3h (Supplementary Methods) (**Fig. 9A**).

**Fig. 9.**
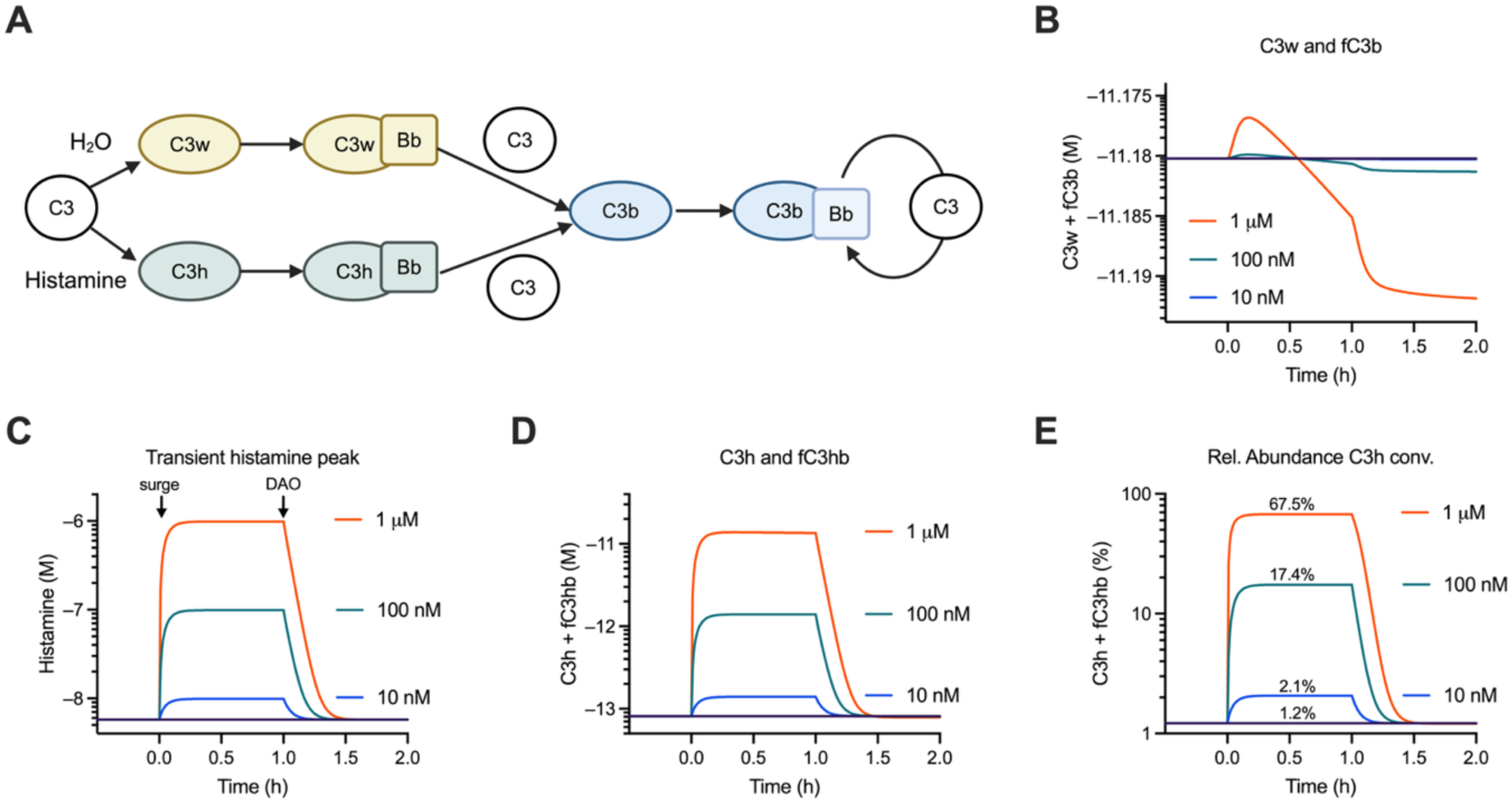
Quantitative modeling of C3h. (**A**) Schematic modeling of histamine-conjugated C3 species (C3h) alongside C3(H_2_O) (C3w). (**B**) Concentration of C3(H_2_O) (C3w) and total fluid-phase C3b, expressed in antilog units, as a function of histamine steady-state concentrations (reference line: 5.8 nM; blue, 10 nM; green, 100 nM; and orange, 1 µM). (**C**) Histamine peak is modeled as a sudden discharge (surge) that is sustained for 1 h until the induction of histamine diamine oxidase (DAO) dampens it. Each curve is characterized by the steady-state concentration of histamine that is reached at its maximum (indicated). Histamine concentration is expressed in antilog units. (**D**) Generation of C3h and fluid-phase C3hb derived from it by various histamine concentration pulses. (**E**) Percentage of C3h and fluid-phase C3hb over the total amount of fluid-phase C3h, C3(H_2_O) (C3w) and C3b generated from every possible route.

C-model simulations performed under physiological histamine concentration showed that C3h-based C3 convertases could reach baseline concentrations in blood (∼1 pM) that are between 1-10% of those of C3 convertases assembled from C3(H_2_O) (**Fig. 9B,C**). The concentration of C3(H_2_O)-based C3 convertase constitutes a useful baseline reference, as it defines the priming level of the complement system (**Fig. 9B**). Modeling the effect of time-limited histamine peak concentrations between 0.6 nM and 1 µM on the abundance of C3h and C3hb showed that C3h-based convertases could equal or even outweigh the overall concentration of C3(H_2_O)-based convertases at and above 100 µM histamine (**Fig. 9D,E**). Quantitative modelling of these processes demonstrates that the transient nature of histamine pulses allows a strong, but local, anaphylactic response to develop and then subside. For patients affected by chronically elevated histamine concentration or harboring genetic defects impairing the control of complement’s alternative pathway activation, we predict a larger amount of C3h-based convertases and a correspondingly greater level of complement activation through the alternative pathway.

These findings suggest that chronic as well as transient histamine levels, among other factors, can influence the baseline (priming) level of complement activation in otherwise healthy individuals, and that this modulation effect of histamine could have a more substantial impact on local and systemic complement activation for patients with either histamine or complement-associated conditions.

## Discussion

In this work, we have described and characterized C3h, a novel species of C3 that is generated by the reaction between the biologically active histamine and the high-energy thioester bond present in the TED domain of complement factor 3 (C3). The small size of this posttranslational modification (∼110 Da) and the vanishingly small amounts that can be produced *in vivo* help to explain why it had not been identified previously, even though other C3-derived products obtained through the reaction of the thioester bond with water or other primary amines are well known.

We have determined the crystal structure of C3h in its C3b-like form, after having lost the C3a domain through spontaneous chemical cleavage. In addition, we have characterized the structure of the complete, mature C3h (including C3a) in solution by SAXS and through extensive MD simulations. The crystal structure has revealed an architecture that is nearly identical to that of C3b, indicating that C3h is a fully active form of C3 in fluid phase. The main difference, the presence of the histamine adduct in Gln1013, does not induce any conformational changes in the TED domain of C3h compared to C3b. In keeping with this observation, C3h can assemble catalytically active C3 convertases through the same biochemical route employed by C3. Furthermore, C3h-containing C3 convertases can be downregulated by FH and FI. As expected from a C3b-like conformation with the CUB and TED fully displaced from their original locations in C3, the cleavage site for FB in C3h becomes solvent-exposed and, therefore, labile to spontaneous hydrolysis or to the action of nonspecific proteases. MD simulations further support the notion that C3a-containing C3h and, by extension, C3(H_2_O), should behave similarly to authentic C3b-containing C3 convertases, as C3a hinders neither the serine proteases responsible for activation (FB and FD) nor the negative regulators like FH and FI.

Our QM/MM studies on the nucleophilic attack of either water or histamine onto the thioester’s carbonyl moiety of Gln1013 provide evidence for an S_N_2 reaction mechanism, whereby the formation of a carbon-oxygen or a carbon-amine bond occurs in a concerted fashion with the breakdown of the thioester bond. Strikingly, the reaction with water is several orders of magnitude more slowly than with histamine (*t*_1/2_ ∼15 d versus 4 s, or a 3 · 10^5^-fold faster reaction), even though the thioester bond sits at the bottom of a very narrow cavity protected from the solvent. The stronger nucleophilic character of the deprotonated amine function and the presence of the imidazole ring are two determinants of the accelerated reaction rate with histamine. These results indicate that while the chemical step of thioester bond hydrolysis could be rate-limiting in the transformation of C3 into C3(H_2_O), the reaction with histamine is several orders of magnitude faster and, therefore, would not be rate-limiting in the transformation from C3 to C3h.

This study demonstrates how easily C3-activated species, such as C3(H_2_O) or C3h, and possibly others, can form *in vivo*. The high energy barrier for C3(H_2_O) is overcome by the ubiquitous presence of water and the high concentration of C3 in blood (∼1 mg/ml, or 5.4 µM). For histamine, in contrast, it is the considerably higher reactivity of the biological amine with the thioester that enables this process. The relative ease with which C3 can be activated keeps complement’s alternative pathway primed and shifts the control of the activation state to the strongly downregulatory activity of FH and FI.

An interesting question concerns the physiological effect of histamine on the activation level of the complement system. In plasma, endogenous histamine is typically present at a concentration of 2.7-9.0 nM. During allergic and anaphylactoid reactions, histamine concentration can increase to greater than 10 nM (90-5000 nM) through the release of histamine granules by mast cells and basophils (*29*). Likewise, other chemically related amines, such as tryptamine, and the derived amines 5-hydroxy tryptophan and serotonin, and phenylethylamine, are also physiologically present in plasma at similar or higher concentrations (3 nM for tryptamine (*30*) and 29 nM for phenylethylamine (*31*)) and in the brain at much lower concentrations, despite playing essential roles in brain chemistry. Histamine and other amines can also be taken up exogenously in the diet in food and drinks. Although a precise toxicological dose for histamine has been challenging to establish, concentration thresholds of 100 mg histamine/kg food and 2 mg/L in alcoholic beverages have been suggested, which would yield upper-bound histamine concentrations in plasma of 900 nM and 15 nM, respectively (assuming standard absorption of 1% into plasma for histamine in healthy individuals). A 2018 meta-analysis of histamine food poisonings concluded that the mean histamine content in poisoning food samples was 1107.21 mg/kg, which would translate into upper-bound histamine concentrations in plasma above 9000 nM. However, these upper-bound concentrations are highly unlikely to be realized due to the rapid turnover of histamine, both extracellularly by diamine oxidase (DAO) and intracellularly by histamine methyltransferase (HMNT).

Individuals with histamine disorders, characterized by sustained elevated levels of histamine, can also exhibit elevated levels of complement activation. This observation has been attributed to the feedback loop between the release of C3a and C5a anaphylatoxins during complement activation and mast cell activation through C3a and C5a binding to their receptors on the surface of mast cells. In mast cell activation syndrome (MCAS) and systemic mastocytosis, the abnormal release of histamine from mast cells has been correlated with the levels of specific complement components, such as C3a and C4a, which can further stimulate mast cells, creating a feedback loop that exacerbates symptoms. During anaphylaxis, allergens crosslinking IgE antibodies on the surface of mast cells and basophils can activate the classical pathway of the complement system, leading to an increased release of C3a and C5a, which can enhance histamine release and contribute to inflammation and anaphylaxis. The crosstalk between histamine and complement revolves around the stimulation of inflammation and the recruitment of immune cells, which sets in motion a feedback loop whereby complement activation increases histamine release, which, in turn, influences the activation and function of complement proteins, thereby perpetuating the cycle of inflammation. In this work, we propose a more direct mode of interaction between histamine and the complement system, based on the observation that histamine can react with the high-energy thioester bond and directly activate complement’s alternative pathway, yielding fluid-phase C3 convertases and releasing C3a (**Fig. 10**).

**Fig. 10.**
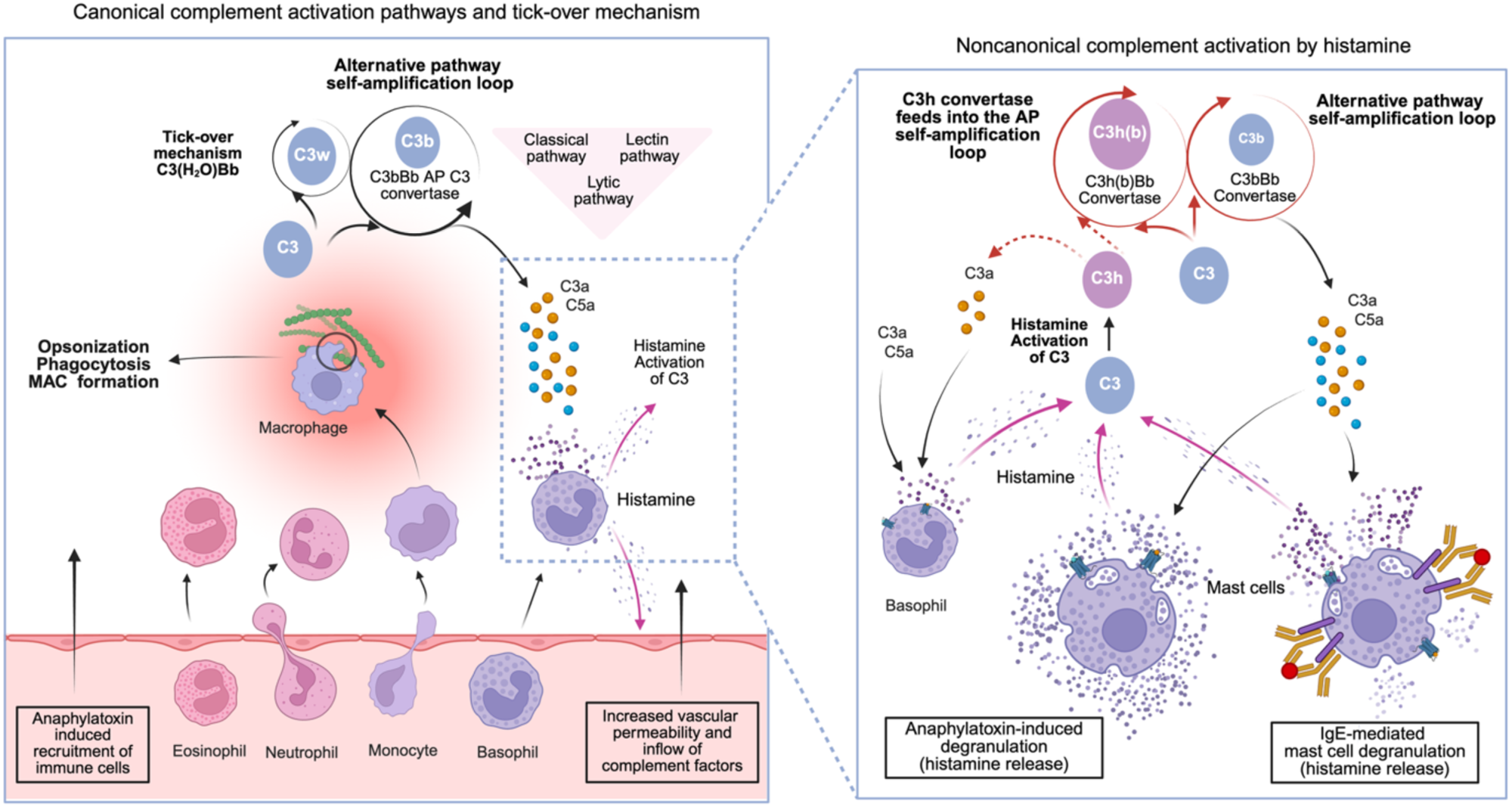
Proposed feedback loop between histamine degranulation and complement activation. The schematic (left) represents the well-established link between the generation of anaphylatoxin chemical gradients during complement activation and increased histamine degranulation. The direct activation of C3 by reaction with histamine is indicated by the inset and magnified on the right. The histaminylation of C3 leads to the assembly of additional C3 convertases that increase the baseline activation level or “priming” of the complement system. By generating C3a and, indirectly, also C5a, C3h(b)Bb C3 convertases close the feedback loop initiated by anaphylatoxin-induced histamine release by granulocytes.

These observations bear clinical, diagnostic, and therapeutic implications. For example, patients with chronic urticaria often have elevated levels of histamine and complement components. Autoantibodies against the high-affinity IgE receptor or IgE itself can activate the complement system, leading to chronic inflammation and histamine release. Hereditary angioedema (HAE), a primary disorder of complement regulation (deficiency of C1 inhibitor), can involve elevated histamine levels during acute attacks, contributing to symptoms such as swelling and pain. In fact, elevated levels of histamine, tryptase (a marker of mast cell activation), and complement components (C3a, C4a, C5a) can be measured to assess the involvement of these pathways in histamine-related disorders. Depending on the results, treatments based on antihistamines or complement inhibitors (such as C1 inhibitors for HAE) can be used to manage symptoms by reducing complement activation. The strong correlation between certain histamine disorders and inflammatory pathologies that activate the complement system suggests that the direct reaction of histamine with C3 may constitute another disease-driving activation pathway.

In conclusion, our study shows that C3-activated species can form in the fluid phase through chemical reaction with histamine (and possibly other biogenic amines), either locally in tissues upon mast cell degranulation or in plasma under conditions of abnormally high histamine concentration. These species, epitomized by C3h, can contribute to the pathophysiology and symptoms of various allergic and inflammatory conditions. While it has been known for years that C3a and C5a can trigger histamine release from mast cells and basophils, our findings indicate that there exists a closed feedback loop whereby histamine can also trigger the low-level activation of C3 (and possibly other thioester-containing proteins, such as a C4), at least to the same level attained by the spontaneous tick-over hydrolysis of C3. Therefore, understanding the relationship between complement activation and histamine might be crucial for effective diagnosis and treatment of a variety of inflammatory and/or complement-associated conditions.

## Materials and Methods

### Generation of C3h

To obtain C3-histamine (C3h), we set up a reaction with 0.5 mg/ml C3 and 0.2 M histamine in 0.1 M sodium phosphate buffer (pH 7.0) and incubated it for 12 h at 4 °C with gentle rotation. The reaction was then desalted over a HiPrep 26/10 Desalting column (Cytiva) equilibrated in 20 mM Tris-HCl (pH 7.5), 100 mM NaCl. To purify C3h from any remaining C3, we loaded the desalted reaction on a Mono Q 5/50 GL column (Cytiva) equilibrated in the same buffer and eluted the bound protein with a 0-100% gradient of 20 mM Tris-HCl (pH 7.5), 500 mM NaCl over 20 column volumes. Finally, peak fractions were pooled and dialyzed against a buffer containing 10 mM HEPES-NaOH (pH 7.4), 150 mM NaCl.

### Mass spectrometry identification of the histamine adduct

A total volume of 1 μl of C3 or C3h, equivalent to 1 μg of total protein, was processed on the day of analysis. The samples were digested with trypsin on S-Trap microcolumns under denaturing conditions. After digestion, salts were removed using Ziptip C18 tips, and the digestion product was quantified using the Qubit method. Subsequently, an equivalent of 500 ng of the digest from each sample (with a theoretical maximum amount of peptide ∼10 ng) was monitored using targeted proteomics in Parallel Reaction Monitoring (PRM) format. The liquid chromatography system used was a Thermo Ultimate 3000 RSLC nano-LC, coupled to a Thermo Orbitrap Exploris OE240 mass spectrometer, configured for targeted scan data acquisition. The analysis focused on the m/z values of 1986.42 and 2032.96, corresponding to the +2 charge state of the peptides ^1002^HLIVTPSGCGEQNMIGMTPTVIAVHYLDETEQWEK^1036^ and the corresponding histamine adduct. Separation was performed on a reversed-phase C18 column with dimensions of 15 cm in length and 75 μm in internal diameter, using a 60-minute elution gradient.

### Kinetics of histamine addition

The rate of histamine reaction was determined by following the titration of Cys1010 free sulfhydryl group with 5,5’-dithiobis(2-nitrobenzoic acid) (DTNB, Ellman’s reagent) after thioester bond aminolysis. Reactions were assembled in 100 µl with 5.4 µM C3, 14 µM DTNB, and 100 mM histamine in 50 mM sodium phosphate buffer (pH 7.4), and the reaction was monitored spectrophotometrically at 412 nm (*25*). For comparison, the same reaction was carried out substituting methylamine for histamine at the same concentration. *A*_412_ values were transformed to sulfhydryl concentration units using the reported extinction coefficient of 1.36 × 10^4^ M^-1^ cm^-1^ and normalized with respect to the total concentration of C3 (*32*). Progress data was fitted to a hyperbola to calculate the kinetic rate of the histamine addition reaction with Prism GraphPad 10.

### Histamine reaction

The effect of histamine concentration on the amount of C3h generated was evaluated. Reactions (20 µl) were set up with 2 µg C3 and varying concentrations of histamine (from 0.1 µM to 0.2 M) and incubated for 4 h at 37 °C. Next, 75 nM FH and 47.5 nM FI were added to the reaction, and incubation was continued for 30 min to convert C3(H_2_O) and C3h (but not C3) into their corresponding iC3b forms. 10 µl were withdrawn, combined with 2 µl of 4X SDS-PAGE gel loading buffer [250 mM Tris, 8% (w/v) SDS, 40% (v/v) glycerol, 0.05 (w/v) bromophenol blue, 4% (v/v) β-mercaptoethanol], and subjected to boiling at 80 °C for 5 min to halt the reaction. Subsequently, reaction products were separated by SDS-PAGE gel electrophoresis on either 10% or 4-20% Bio-Rad precast acrylamide gels, followed by staining with Coomassie Brilliant Blue (CBB) or Western blotting. Stained gels were then digitized using a ChemiDoc Touch Imaging System (Bio-Rad). Quantification of the extent of cleavage was performed by analyzing the histamine concentration-dependent disappearance of the α’ chain of C3b or the α chain of C3h. This involved band quantification using ImageJ software (https://imagej.net/ij/), complemented by fitting the data to a 4-parameter log-logistic function with Prism GraphPad 10.

### FH-mediated cleavage by FI

A time course analysis of FH-mediated cleavage by FI of C3b or C3h was conducted under controlled conditions. The reaction was set up in a thermostated water bath at 37 °C, comprising 0.92 µM C3b or C3h, 75 nM FH, and 47.5 nM FI (added last) in a reaction buffer consisting of 10 mM HEPES-NaOH (pH 7.4) and 150 mM NaCl. At designated time points (0-60 min), 6 µl aliquots of the reaction mixture (containing 1 µg C3b/C3h, 70 ng FH, and 25 ng FI) were withdrawn, combined with 2 µl of 4X SDS-PAGE gel loading buffer [250 mM Tris, 8% (w/v) SDS, 40% (v/v) glycerol, 0.05 (w/v) bromophenol blue, 4% (v/v) β-mercaptoethanol], and subjected to boiling at 80 °C for 5 min to halt the reaction. Subsequently, reaction products were separated by SDS-PAGE gel electrophoresis on either 10% or 4-20% Bio-Rad precast acrylamide gels, followed by staining with Coomassie Brilliant Blue (CBB). Stained gels were then digitized using a ChemiDoc Touch Imaging System (Bio-Rad). Quantification of the extent of cleavage was performed by analyzing the time-dependent disappearance of the α’ chain of C3b or the α chain of C3h. This involved band quantification using ImageJ software (https://imagej.net/ij/), complemented by fitting the data to a 4-parameter log-logistic function using SciPy (https://scipy.org).

### Trypsin susceptibility assay

A limited proteolysis assay was established to evaluate the increased susceptibility of C3h to trypsin cleavage compared to C3. Initially, we determined experimentally the optimal mass ratio of trypsin to C3 or C3h required to digest 50% of 1.5 µg of the substrate protein within 5 min at room temperature (25 °C), as assessed by SDS-PAGE electrophoresis on 10% acrylamide gels. Various mass ratios ranging from 1:10 to 1:10^6^ were tested, and the optimal mass ratio was determined to be 1:100. Subsequently, for the optimized trypsin susceptibility assay, 22.5 µg of C3 or C3h and 0.225 µg of trypsin (at a 1:100 mass ratio) were mixed in 180 µl of reaction buffer containing 10 mM HEPES-NaOH (pH 7.4) and 150 mM NaCl. At designated time points (0-60 minutes), 12 µl aliquots were withdrawn, and the reaction was stopped by mixing with 4 µl of 4X SDS-PAGE gel loading buffer [250 mM Tris, 8% (w/v) SDS, 40% (v/v) glycerol, 0.05% (w/v) bromophenol blue, 4% (v/v) β-mercaptoethanol] and boiling at 80 °C for 5 min. Subsequently, the reaction products were separated by SDS-PAGE gel electrophoresis on either 10% or 4-20% Bio-Rad precast acrylamide gels, followed by staining with Coomassie Brilliant Blue (CBB). Stained gels were then digitized using a ChemiDoc Touch Imaging System (Bio-Rad). Quantification of the extent of cleavage was performed by analyzing the time-dependent disappearance of the α’ chain of C3b or the α chain of C3h. This involved band quantification using ImageJ software (https://imagej.net/ij/), supplemented by fitting the data to an exponential decay function using SciPy (https://scipy.org).

### Hemolytic assays

The capacity of C3h to promote complement activation on cellular surfaces was assessed in hemolytic assays using guinea pig erythrocytes (GpEs). In brief, GpEs (0.5% packed cell volume; TCS Biosciences) in AP buffer, consisting of veronal buffer saline (2.5 mM barbital, 1.5 mM sodium barbital, 144 mM sodium chloride, pH 7.4) with 5 mM magnesium chloride, 8 mM EGTA, and 0.1% gelatin, were incubated with 10% normal human serum (NHS) in AP buffer and increasing amounts of C3h (0–5%) for 30 min at 37 °C. Reactions were stopped using AP buffer containing 20 mM EDTA. After centrifugation, supernatants were read at 414 nm. Erythrocytes diluted in AP buffer were used as blanks for spontaneous lysis, and serum without added FH was considered the reference for 100% lysis. To set the experimental conditions in which increasing amounts of C3h were analyzed, we chose the minimal concentration of FH resulting in 20% lysis. Data was plotted and analyzed with Prism GraphPad 10.

### X-ray crystallography

C3h was adjusted to 5 mg/ml in 10 mM HEPES-NaOH (pH 7.4), 150 mM NaCl buffer. Sitting-drop vapor-diffusion crystallization experiments were done at 20 °C with drops consisting of 1 μl of protein complex plus 1 μl reservoir solution [10% (w/v) PEG-MME 2000, 0.1 M sodium acetate, 0.1 M Bis-Tris propane (pH 7.8)]. Crystals were collected, soaked in a reservoir solution containing 15% (v/v) sterile glycerol as cryoprotectant, flash-cooled, and stored in liquid nitrogen until data collection. The X-ray diffraction data extended to 2.81-Å resolution. Diffraction data were collected at cryogenic temperature (100 *K*) at the BL13-XALOC beamline (ALBA Synchrotron, Barcelona, Spain) (*33*). The diffraction data were processed with XDS/SCALE (*34*). The structure of C3h was solved by molecular replacement with *PHASER* (*35*) using the structure of C3b from PDB 5FO7 as a search model (*36*). Iterative model building and refinement were performed in *Coot* (*37*) and *PHENIX* (*38*), respectively, and validation in MolProbity (*39*).

The coordinates and structure factors of C3h have been deposited with PDB accession code 9SKN.

### In-line size-exclusion chromatography coupled SAXS (SEC-SAXS)

Synchrotron SAXS measurements [*I*(*q*) vs. *q*, where *q* = 4 π sinθ/λ, 2θ is the scattering angle, and λ = 0.9464 Å] were performed in SEC-SAXS mode at 15 °C at the B21 beamline at the Diamond Light Source (Didcot, UK) (*40*). **Supplementary Table S1** summarizes the SAXS data and provides additional information. 50 µl of 1.2 mg/ml C3h was injected into a Shodex KW-403 chromatographic column (separation range between 10-700 kDa) equilibrated in 20 mM Tris-HCl (pH 8.0), 150 mM NaCl, 3% (w/v) glycerol, and separated at a flow rate of 160 µl/min on a dual Agilent 1260 HPLC system. SAXS measurements collected across the main chromatographic peak were averaged and buffer subtracted to produce a reliable measurement of C3h. In this way, SEC-SAXS achieves the purification of the protein sample during SAXS measurements, improving buffer matching and reducing polydispersity. BioXTAS RAW (*41*) and ATSAS 3.2 (*42*) were used to process the scattering data, extract structural information, perform *ab initio* shape restoration, and rigid-body fitting against C3h scattering profile. The extrapolated forward scattering at zero concentration, *I*(0), and the radius of gyration, *R*_g_, were calculated from the Guinier approximation and from the real-space pair-distance distribution function (*P*(*r*) vs. *r*) calculated with GNOM (*43*). From the *P*(*r*) profile, the maximum particle dimension (*D*_max_) could be evaluated. Concentration-independent methods were used to estimate the molecular mass of C3h (*44*). C3h shape was restored with DAMMIF (*45*). The final average model of C3h has an estimated resolution of ∼31 Å with SASRES (*46*). The agreement between the SAXS scattering profile and the structural models was assessed with CRYSOL (*47*).

### Homology modeling

We used the homology modeling software MODELLER (*48*) to build complete models of C3h, including the C3a anaphylatoxin domain, starting from the composite model obtained by SAXS-based rigid-body fitting with SASREF (*49*), and C3, starting from a previously determined structure (PDB 2A73) (*50*). A loop modeling and refinement procedure (*51*) was deployed to fill in the following gaps in the structure: residues 76-77, 727-729, and 1351-1358 (C3h) and residues 720-728 and 1107-1112 (C3). All (5) output models were ranked according to the DOPE score (*52*) (C3h and C3), and their agreement with the experimental SAXS scattering profile was assessed with CRYSOL (*47*) (C3h).

### Molecular dynamics

All molecular dynamics (MD) simulations were carried out with GROMACS v. 2023 in the Galician Supercomputing Center (CESGA). A molecular system consisting of C3(H_2_O) (to mimic C3h) was constructed and minimized using GROMACS (*53*). The initial model for C3(H_2_O) was obtained from C3h with MODELLER (*48*). The OPLS-AA/L forcefield and SPC/E water model were used (*54*). Solvent counterions were added to neutralize the system’s net charge at a physiological concentration (135 mM NaCl). The systems were first subjected to minimization, followed by 1.25-ns equilibration of the NVT and NPT ensembles. All production runs (1.1 µs in triplicate each) were carried out in the NPT ensemble at 300 K and 1 bar. Temperature was controlled by Nosé−Hoover (*55*, *56*) (coupling constant τ*_t_* = 1.0 ps) and pressure by Parrinello−Rahman (*57*, *58*) (τ*_p_* = 1.0 ps) schemes. Periodic boundary conditions were applied in three-dimensional space, and electrostatic forces were calculated with the Particle Mesh Ewald (PME) method (*59*, *60*) using a real-space cutoff of 1.2 nm. Equations of motion were integrated using the leapfrog scheme with a time step of 2 fs. The last 1 µs of the trajectories were analyzed, and the structures were clustered using the GROMOS algorithm (*61*). For C3(H_2_O), protein structural snapshots (1 ns/snapshot) were evaluated as to their agreement with the SAXS scattering profile for C3h using CRYSOL (*47*).

### QM/MM and MM Free Energy Calculations

Reaction mechanisms of the thioester bond with water and histamine were carried out at the QM/MM level. The Adaptive String Method (ASM) (*62*) was used to characterize the free energy landscape of the chemical reactions. This identifies the minimum free energy path (MFEP) on a multidimensional free energy surface, defined by a set of collective variables (CVs) that incorporate all the distances associated with bonds being formed or broken during the process.

The full system was divided into a QM and an MM region. For the aminolysis reaction, the QM region included the side chains of Cys1010 and Glu1013 (as the Gln1013 side chain amide group is lost upon thioester bond formation) and the full histamine molecule. For the hydrolysis process, the QM region is composed of the side chains of Cys1010 and Glu1013, the nucleophilic water molecule, and the side chain of Glu1012, the glutamate residue immediately preceding Gln1013 in the primary sequence of C3. All other atoms were treated at the MM level with the ff14SB force field for the protein (*63*) and TIP3P for water (*64*). The QM region was described using the B3LYP functional (*65*, *66*) with a 6-31+G* basis set and D3 dispersion corrections (*67*). All QM/MM calculations were performed using a modified version of Amber18 (*68*) coupled with Gaussian16 (*69*) for density functional theory calculations. A cut-off radius of 15 Å was used for all QM/MM interactions.

To explore the mechanistic proposals, 96 replicas of the system (string nodes) connected the reactant and product structures in a space defined by the CVs shown in **Supplementary Fig. S5 and S6**. These nodes were moved according to their free energy gradient obtained by means of QM/MM MD simulations carried out at 300 *K* using Langevin dynamics with a collision frequency of 5.0 ps^-1^. The nodes are redistributed equidistantly along the string to avoid falling into the global minima region (reactants and products). This process was continued until the string converged to the MFEP with an RMSD for the CVs below 0.1 amu^1/2^·Å for at least 2 ps. Replica exchanges between nodes were attempted every 50 steps to ensure convergence. Upon convergence, a path-CV (*70*) measuring the progress of the system along the MFEP from reactants to products was defined and used to trace the free energy profile of the reaction using umbrella sampling (*71*). For this purpose, each node was subjected to 10 ps of QM/MM simulations using WHAM as the integration method (*72*). To maintain a homogeneous probability density distribution of the reaction coordinate, the force constants used to bias the ASM simulations were determined on the fly. The integration time for the QM/MM simulations was set to 1 fs and the mass of the transferred hydrogen atoms was set to 2 amu.

To evaluate the free energy difference between monocationic and free base forms of histamine (1′G_his_) in the active site of C3h at a given pH we used the following thermodynamic relationship:

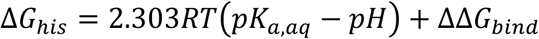

where the pH was set equal to 7.4, the p*K*_a_ of the terminal amino group in aqueous solution was taken as 9.76 (*26*) and ΔΔG_bind_ is the free energy change in the binding free energy of the protonated and free base forms of histamine in C3h. This free energy change was evaluated at the MM level from the free energy differences resulting in the alchemical transformations of the free base histamine into the monocationic form both in aqueous solution and in the protein environment (**Supplementary Fig. S7**).

Simulations were carried out using Amber18 with the ff14SB and TIP3P force fields. Free energy calculations were performed using the dual topology approach and following a published protocol for the thermodynamic integration method (*73*). Briefly, the temperature was kept at 300 *K* using Langevin dynamics with a collision frequency of 5.0 ps^-1^. The pressure was kept constant at 1.01325 bar with Monte Carlo barostat and a pressure relaxation time of 2.0 ps. The atom that disappears during the alchemical transformation (a proton) was assigned to the soft-core region for both van der Waals and electrostatic interactions. All simulations were run in pmemd.cuda in GPU, and the time step was set to 1 fs. In each of the five replicas used to obtain a reliable averaged estimation (**Supplementary Table S2**), the alchemical transformation was performed using nine different λ values, following the Gaussian quadrature method (λ = 0.01592, 0.08198, 0.19331, 0.33787, 0.5, 0.66213, 0.80669, 0.91802, and 0.98408). The five separate replicas were run for 5 ns for each λ value. The first 1 ns of each simulation was considered an equilibration step at the corresponding λ window and was therefore skipped for thermodynamic integration. The average value of the free energy change corresponding to the alchemical transformation of the residue in water (–44.83 ± 0.04 kcal·mol^-1^) was then subtracted from the average value obtained in the protein environment (–44.58 ± 0.58 kcal·mol^-1^).

### Quantitative simulation of the complement system

We performed quantitative simulations of complement activation biomarkers using the complement-specific enhanced pharmacokinetics simulation environment C-model (*28*), after incorporating all reactions involving histamine, C3h, and C3h-derived proteolytic and hydrolytic fragments (see Supplementary Methods for the implementation details). C-model incorporates all three complement activation pathways and can be adapted to simulate various pathophysiological scenarios.

## Supporting information

Main Text

## Acknowledgments

The authors acknowledge the ALBA synchrotron light source (Barcelona, Spain) for providing synchrotron radiation facilities at the BL13-XALOC beamline, the Diamond Light Source (Didcot, UK) for access to the BioSAXS ID29 beamline, and the Galicia Supercomputing Center (CESGA) for providing computational support. We also acknowledge the CSIC Network of Rare Diseases (RER-CSIC) (MCV). KPG and LAG acknowledge the support of the PhD program in Biochemistry, Molecular Biology, and Biomedicine, and the PhD program in Medicinal Chemistry, respectively, at the Universidad Complutense de Madrid (UCM).

## Author contributions

Conceptualization: FJF, SRC, IT, MCV

Methodology: KPG, JQG, IMM, HMM, LAG, CAR, FJF, SRC, IT, MCV

Investigation: HMM, SRC, CAR, IT, FJF, LAG, MCV

Visualization: CAR, IT, FJF, LAG, MCV

Supervision: FJF, IT, SRC, MCV

Writing—original draft: FJF, IT, MCV

Writing—review & editing: FJF, IT, SRC, MCV

## Data and materials availability

All data needed to evaluate the conclusions in the paper are present in the paper and/or the Supplementary Materials. Coordinate and structure factor files for C3h have been deposited in the Protein Data Bank under PDB accession code 9SKN.

## References

1. M. J. Walport, Complement. First of two parts. N. Engl. J. Med. 344, 1058–1066 (2001).

2. M. J. Walport, Complement. Second of two parts. N. Engl. J. Med. 344, 1140–1144 (2001).

3. F. J. Fernández, S. Gómez, M. C. Vega, Pathogens’ toolbox to manipulate human complement. Semin. Cell Dev. Biol. 85, 98–109 (2019).

4. B. V. Geisbrecht, J. D. Lambris, P. Gros, Complement component C3: A structural perspective and potential therapeutic implications. Semin. Immunol. 59, 101627 (2022).

5. S. K. A. Law, A. W. Dodds, The internal thioester and the covalent binding properties of the complement proteins C3 and C4. Protein Sci. 6, 263–274 (1997).

6. U. Amara, M. A. Flierl, D. Rittirsch, A. Klos, H. Chen, B. Acker, U. B. Brückner, B. Nilsson, F. Gebhard, J. D. Lambris, M. Huber-Lang, Molecular Intercommunication between the Complement and Coagulation Systems. J. Immunol. 185, 5628–5636 (2010).

7. D. Ricklin, G. Hajishengallis, K. Yang, J. D. Lambris, Complement: a key system for immune surveillance and homeostasis. Nat. Immunol. 11, 785–797 (2010).

8. H. I. Kenawy, I. Boral, A. Bevington, Complement-coagulation cross-talk: a potential mediator of the physiological activation of complement by low pH. Front. Immunol. 6 (2015).

9. B. Nilsson, K. Nilsson Ekdahl, The tick-over theory revisited: Is C3 a contact-activated protein? Immunobiology 217, 1106–1110 (2012).

10. M. E. Parsons, C. R. Ganellin, Histamine and its receptors. Br. J. Pharmacol. 147 (2006).

11. P. Lieberman, The basics of histamine biology. Ann. Allergy. Asthma. Immunol. 106, S2–S5 (2011).

12. J. Dyer, K. Warren, S. Merlin, D. D. Metcalfe, M. Kaliner, Measurement of plasma histamine: description of an improved method and normal values. J. Allergy Clin. Immunol. 70, 82–87 (1982).

13. K. Kimura, M. Adachi, K. Kubo, H1- and H2-receptor antagonists prevent histamine release in allergic patients after the administration of midazolam-ketamine. A randomized controlled study. Inflamm. Res. 48, 128–132 (1999).

14. E. Costing, E. Neugebauer, J. J. Keyzer, W. Lorenz, Determination of histamine in human plasma: the European external quality control study 1988. Clin. Exp. Allergy 20, 349–357 (1990).

15. V. Zdravkovic, S. Pantovic, G. Rosic, A. Tomic-Lucic, N. Zdravkovic, M. Colic, Z. Obradovic, M. Rosic, Histamine blood concentration in ischemic heart disease patients. J. Biomed. Biotechnol. 2011, 315709 (2011).

16. M. M. Dale, J. C. Foreman, “Histamine as a mediator of allergic and inflammatory reactions” in Textbook of Immunopharmacology (Blackwell Scientific Publications, Boston, MA, ed. 3rd, 1993), pp. xiii, 370.

17. M. Dy, E. Schneider, Histamine–cytokine connection in immunity and hematopoiesis. Cytokine Growth Factor Rev. 15, 393–410 (2004).

18. N. R. Cooper, G. R. Nemerow, The role of antibody and complement in the control of viral infections. J. Invest. Dermatol. 83, 121s–127s (1984).

19. Y. Yanase, Y. Matsuo, S. Takahagi, T. Kawaguchi, K. Uchida, K. Ishii, A. Tanaka, D. Matsubara, K. Ozawa, M. Hide, Coagulation factors induce human skin mast cell and basophil degranulation via activation of complement 5 and the C5a receptor. J. Allergy Clin. Immunol. 147, 1101–1104.e7 (2021).

20. D. Lappin, H. L. Moseley, K. Whaley, Effect of histamine on monocyte complement production. II. Modulation of protein secretion, degradation and synthesis. Clin. Exp. Immunol. 42, 515–522 (1980).

21. A. Falus, E. Walcz, M. Brozik, H. Rokita, G. Fust, A. Hajnal, K. Meretey, Stimulation of histamine receptors of human monocytoid and hepatoma-derived cell lines and mouse hepatocytes modulates the production of the complement components C3, C4, factor B, and C2. Scand. J. Immunol. 30, 241–248 (1989).

22. P. Sánchez-Corral, L. C. Antón, J. M. Alcolea, G. Marqués, A. Sánchez, F. Vivanco, Separation of active and inactive forms of the third component of human complement, C3, by fast protein liquid chromatography (FPLC). J. Immunol. Methods 122, 105–113 (1989).

23. 23. M. M. Ruseva, M. Heurich, “Purification and characterization of human and mouse complement C3” in The Complement System: Methods and Protocols, M. Gadjeva, Ed. (Humana Press, Totowa, NJ, 2014; 10.1007/978-1-62703-724-2_6), pp. 75–91.

24. B. J. C. Janssen, A. Christodoulidou, A. McCarthy, J. D. Lambris, P. Gros, Structure of C3b reveals conformational changes that underlie complement activity. Nature 444, 213–216 (2006).

25. M. K. Pangburn, R. D. Schreiber, H. J. Müller-Eberhard, Formation of the initial C3 convertase of the alternative complement pathway. Acquisition of C3b-like activities by spontaneous hydrolysis of the putative thioester in native C3. J. Exp. Med. 154, 856–867 (1981).

26. L. Settimo, K. Bellman, R. M. A. Knegtel, Comparison of the accuracy of experimental and predicted pKa values of basic and acidic Compounds. Pharm. Res. 31, 1082–1095 (2014).

27. J. Wu, Y.-Q. Wu, D. Ricklin, B. J. C. Janssen, J. D. Lambris, P. Gros, Structure of complement fragment C3b–factor H and implications for host protection by complement regulators. Nat. Immunol. 10, 728–733 (2009).

28. L. Alfonso-González, M. C. Vega, F. J. Fernández, C-model: A comprehensive enhanced pharmacokinetic/pharmacodynamic simulation environment targeting the complement system. British Journal of Pharmacology (2025) 1:19. 10.1111/bph.70054. *Br. J. Pharmacol.* **1**, 19 (2025).

29. D. Laroche, M.-C. Vergnaud, B. Sillard, H. Soufarapis, H. Bricard, Biochemical Markers of Anaphylactoid Reactions to Drugs Comparison of Plasma Histamine and Tryptase. Anesthesiology 75, 945–949 (1991).

30. L. Palego, L. Betti, A. Rossi, G. Giannaccini, Tryptophan Biochemistry: Structural, Nutritional, Metabolic, and Medical Aspects in Humans. J. Amino Acids 2016, 8952520 (2016).

31. M. Tsuji, K. Ohi, C. Taga, T. Myojin, S. Takahashi, Determination of β-phenylethylamine concentrations in human plasma, platelets, and urine and in animal tissues by high-performance liquid chromatography with fluorometric detection. Anal. Biochem. 153, 116–120 (1986).

32. G. L. Ellman, Tissue sulfhydryl groups. Arch. Biochem. Biophys. 82, 70–77 (1959).

33. J. Juanhuix, F. Gil-Ortiz, G. Cuní, C. Colldelram, J. Nicolás, J. Lidón, E. Boter, C. Ruget, S. Ferrer, J. Benach, Developments in optics and performance at BL13-XALOC, the macromolecular crystallography beamline at the Alba Synchrotron. J. Synchrotron Radiat. 21, 679–689 (2014).

34. W. Kabsch, XDS. Acta Crystallogr. D Biol. Crystallogr. 66, 125–132 (2010).

35. A. J. McCoy, R. W. Grosse-Kunstleve, P. D. Adams, M. D. Winn, L. C. Storoni, R. J. Read, Phaser crystallographic software. J. Appl. Crystallogr. 40, 658–674 (2007).

36. F. Forneris, J. Wu, X. Xue, D. Ricklin, Z. Lin, G. Sfyroera, A. Tzekou, E. Volokhina, J. C. Granneman, R. Hauhart, P. Bertram, M. K. Liszewski, J. P. Atkinson, J. D. Lambris, P. Gros, Regulators of complement activity mediate inhibitory mechanisms through a common C3b-binding mode. EMBO J. 35, 1133–1149 (2016).

37. P. Emsley, B. Lohkamp, W. G. Scott, K. Cowtan, Features and development of Coot. Acta Crystallogr. Sect Biol. Crystallogr. 66, 486–501 (2010).

38. D. Liebschner, P. V. Afonine, M. L. Baker, G. Bunkóczi, V. B. Chen, T. I. Croll, B. Hintze, L. W. Hung, S. Jain, A. J. McCoy, N. W. Moriarty, R. D. Oeffner, B. K. Poon, M. G. Prisant, R. J. Read, J. S. Richardson, D. C. Richardson, M. D. Sammito, O. V. Sobolev, D. H. Stockwell, T. C. Terwilliger, A. G. Urzhumtsev, L. L. Videau, C. J. Williams, P. D. Adams, Macromolecular structure determination using X-rays, neutrons and electrons: recent developments in Phenix. Acta Crystallogr. Sect. Struct. Biol. 75, 861–877 (2019).

39. V. B. Chen, W. B. Arendall, J. J. Headd, D. A. Keedy, R. M. Immormino, G. J. Kapral, L. W. Murray, J. S. Richardson, D. C. Richardson, MolProbity: all-atom structure validation for macromolecular crystallography. Acta Crystallogr. Sect Biol. Crystallogr. 66, 12–21 (2010).

40. N. P. Cowieson, C. J. C. Edwards-Gayle, K. Inoue, N. S. Khunti, J. Doutch, E. Williams, S. Daniels, G. Preece, N. A. Krumpa, J. P. Sutter, M. D. Tully, N. J. Terrill, R. P. Rambo, Beamline B21: high-throughput small-angle X-ray scattering at Diamond Light Source. J. Synchrotron Radiat. 27, 1438–1446 (2020).

41. J. B. Hopkins, R. E. Gillilan, S. Skou, BioXTAS RAW: improvements to a free open-source program for small-angle X-ray scattering data reduction and analysis. J. Appl. Crystallogr. 50, 1545–1553 (2017).

42. K. Manalastas-Cantos, P. V. Konarev, N. R. Hajizadeh, A. G. Kikhney, M. V. Petoukhov, D. S. Molodenskiy, A. Panjkovich, H. D. T. Mertens, A. Gruzinov, C. Borges, C. M. Jeffries, D. I. Svergun, D. Franke, *ATSAS 3.0*: expanded functionality and new tools for small-angle scattering data analysis. J. Appl. Crystallogr. 54, 343–355 (2021).

43. D. I. Svergun, Determination of the regularization parameter in indirect-transform methods using perceptual criteria. J. Appl. Crystallogr. 25, 495–503 (1992).

44. R. P. Rambo, J. A. Tainer, Accurate assessment of mass, models and resolution by small-angle scattering. Nature 496, 477–481 (2013).

45. D. Franke, D. I. Svergun, DAMMIF, a program for rapid ab-initio shape determination in small-angle scattering. J. Appl. Crystallogr. 42, 342–346 (2009).

46. A. T. Tuukkanen, G. J. Kleywegt, D. I. Svergun, Resolution of *ab initio* shapes determined from small-angle scattering. IUCrJ 3, 440–447 (2016).

47. D. Svergun, C. Barberato, M. H. J. Koch, *CRYSOL* – a program to evaluate X-ray solution scattering of biological macromolecules from atomic coordinates. J. Appl. Crystallogr. 28, 768–773 (1995).

48. A. Fiser, A. Sali, Modeller: generation and refinement of homology-based protein structure models. Methods Enzymol. 374, 461–491 (2003).

49. M. V. Petoukhov, D. I. Svergun, Global rigid body modeling of macromolecular complexes against small-angle scattering data. Biophys. J. 89, 1237–1250 (2005).

50. B. J. C. Janssen, E. G. Huizinga, H. C. A. Raaijmakers, A. Roos, M. R. Daha, K. Nilsson-Ekdahl, B. Nilsson, P. Gros, Structures of complement component C3 provide insights into the function and evolution of immunity. Nature 437, 505–511 (2005).

51. A. Fiser, R. K. Do, A. Sali, Modeling of loops in protein structures. Protein Sci. 9, 1753–1773 (2000).

52. M. Shen, A. Sali, Statistical potential for assessment and prediction of protein structures. Protein Sci. 15, 2507–2524 (2006).

53. M. J. Abraham, T. Murtola, R. Schulz, S. Páll, J. C. Smith, B. Hess, E. Lindahl, GROMACS: High performance molecular simulations through multi-level parallelism from laptops to supercomputers. SoftwareX 1-2, 19–25 (2015).

54. W. L. Jorgensen, D. S. Maxwell, J. Tirado-Rives, Development and testing of the OPLS All-Atom force field on conformational energetics and properties of organic liquids. J. Am. Chem. Soc. 118, 11225–11236 (1996).

55. S. Nosé, A molecular dynamics method for simulations in the canonical ensemble. Mol. Phys. 52, 255–268 (1984).

56. W. G. Hoover, Canonical dynamics: Equilibrium phase-space distributions. Phys. Rev. A 31, 1695–1697 (1985).

57. M. Parrinello, Polymorphic transitions in single crystals: A new molecular dynamics method. J. Appl. Phys. 52, 7182 (1981).

58. S. Nosé, M. L. Klein, Constant pressure molecular dynamics for molecular systems. Mol. Phys. 50, 1055–1076 (1983).

59. T. Darden, D. York, L. Pedersen, Particle mesh Ewald: An N⋅log(N) method for Ewald sums in large systems. J. Chem. Phys. 98, 10089–10092 (1993).

60. U. Essmann, L. Perera, M. L. Berkowitz, T. Darden, H. Lee, L. G. Pedersen, A smooth particle mesh Ewald method. J. Chem. Phys. 103, 8577–8593 (1995).

61. X. Daura, K. Gademann, B. Jaun, D. Seebach, W. F. van Gunsteren, A. E. Mark, Peptide folding: When simulation meets experiment. Angew. Chem. Int. Ed. 38, 236–240 (1999).

62. K. Zinovjev, I. Tuñón, Adaptive finite temperature string method in collective variables. J. Phys. Chem. A 121, 9764–9772 (2017).

63. J. A. Maier, C. Martinez, K. Kasavajhala, L. Wickstrom, K. E. Hauser, C. Simmerling, ff14SB: improving the accuracy of protein side chain and backbone parameters from ff99SB. J. Chem. Theory Comput. 11, 3696–3713 (2015).

64. W. L. Jorgensen, J. Chandrasekhar, J. D. Madura, R. W. Impey, M. L. Klein, Comparison of simple potential functions for simulating liquid water. J Chem Phys 79, 926 (1983).

65. A. D. Becke, Density-functional thermochemistry. III. The role of exact exchange. J. Chem. Phys. 98, 5648–5652 (1993).

66. C. Lee, W. Yang, R. G. Parr, Development of the Colle-Salvetti correlation-energy formula into a functional of the electron density. Phys. Rev. B 37, 785–789 (1988).

67. S. Grimme, J. Antony, S. Ehrlich, H. Krieg, A consistent and accurate *ab initio* parametrization of density functional dispersion correction (DFT-D) for the 94 elements H-Pu. J. Chem. Phys. 132, 154104 (2010).

68. 68. D. A. Case, I.Y. Ben-Shalom, S.R. Brozell, D.S. Cerutti, T. E. Cheatham, V.W.D. Cruzeiro, T.A. Darden, R.E. Duke, D. Ghoreishi, M.K. Gilson, H. Gohlke, A.W. Goetz, D. Greene, R. C. Harris, N. Homeyer, Yandong Huang, S. Izadi, A. Kovalenko, T. Kurtzman, T.S. Lee, S. LeGrand, P. Li, C. Lin, J. Liu, T. Luchko, R. Luo, D.J. Mermelstein, K.M. Merz, Y. Miao, G. Monard, C. Nguyen, H. Nguyen, I. Omelyan, A. Onufriev, F. Pan, R. Qi, D.R. Roe, A. Roitberg, C. Sagui, S. Schott-Verdugo, J. Shen, C.L. Simmerling, J. Smith, R. Salomon-Ferrer, J. Swails, R.C. Walker, J. Wang, H. Wei, R.M. Wolf, X. Wu, L. Xiao, D.M. York, P.A. Kollman, Amber 2018, Unpublished (2018); 10.13140/RG.2.2.31525.68321.

69. M. J. Frisch, G. W. Trucks, H. B. Schlegel, G. E. Scuseria, M. A. Robb, J. R. Cheeseman, G. Scalmani, V. Barone, G. A. Petersson, H. Nakatsuji, X. Li, M. Caricato, A. V. Marenich, J. Bloino, B. G. Janesko, R. Gomperts, B. Mennucci, H. P. Hratchian, J. V. Ortiz, A. F. Izmaylov, J. L. Sonnenberg, Williams, F. Ding, F. Lipparini, F. Egidi, J. Goings, B. Peng, A. Petrone, T. Henderson, D. Ranasinghe, V. G. Zakrzewski, J. Gao, N. Rega, G. Zheng, W. Liang, M. Hada, M. Ehara, K. Toyota, R. Fukuda, J. Hasegawa, M. Ishida, T. Nakajima, Y. Honda, O. Kitao, H. Nakai, T. Vreven, K. Throssell, J. A. Montgomery Jr., J. E. Peralta, F. Ogliaro, M. J. Bearpark, J. J. Heyd, E. N. Brothers, K. N. Kudin, V. N. Staroverov, T. A. Keith, R. Kobayashi, J. Normand, K. Raghavachari, A. P. Rendell, J. C. Burant, S. S. Iyengar, J. Tomasi, M. Cossi, J. M. Millam, M. Klene, C. Adamo, R. Cammi, J. W. Ochterski, R. L. Martin, K. Morokuma, O. Farkas, J. B. Foresman, D. J. Fox, Gaussian 16 Rev. C.01, (2016).

70. K. Zinovjev, I. Tuñón, Exploring chemical reactivity of complex systems with path-based coordinates: Role of the distance metric. J. Comput. Chem. 35, 1672–1681 (2014).

71. G. M. Torrie, J. P. Valleau, Nonphysical sampling distributions in Monte Carlo free-energy estimation: Umbrella sampling. J. Comput. Phys. 23, 187–199 (1977).

72. S. Kumar, J. M. Rosenberg, D. Bouzida, R. H. Swendsen, P. A. Kollman, The weighted histogram analysis method for free-energy calculations on biomolecules. I. The method. J. Comput. Chem. 13, 1011–1021 (1992).

73. X. He, S. Liu, T.-S. Lee, B. Ji, V. H. Man, D. M. York, J. Wang, Fast, accurate, and reliable protocols for routine calculations of protein-ligand binding affinities in drug design projects using AMBER GPU-TI with ff14SB/GAFF. ACS Omega 5, 4611–4619 (2020).

