## Supplementary material for "Molecular basis of noncanonical complement C3 activation by histamine": Main Text

**This PDF file includes:**

Supplementary Text
Figs. S1 to S7
Tables S1 to S2
References (1 to 5)

**Supplementary Text**

C-model implementation of histamine effects on complement activation

The software code for C-model, a mechanistic model of the complement system, was modified to incorporate the effects of histamine on complement system activation based on the findings presented in the main text.

For histamine synthesis and degradation under steady-state conditions, we used a kinetic degradation constant ($k_d$) of $5 \times 10^{-3}$ $s^{-1}$, calculated from its half-life (*1*). The kinetic synthesis constant ($k_s$) was calculated using the formula: $k_s = k_d \times$ [physiological blood concentration of histamine]. The physiological histamine concentration used in the steady-state simulation was 5.83 nM (*2*). Additionally, we considered the role of diamine oxidase (DAO) in histamine degradation, using a steady-state blood concentration of 8.2 pM (*3*) and a $k_d$ of $2.14 \times 10^{-5}$ $s^{-1}$, derived from its half-life (*4*). Histamine degradation by diamine oxidase was modeled using a Michaelis-Menten constant ($K_M$) of 2.8 μM and a catalytic rate constant ($k_{cat}$) of 2.32 $s^{-1}$ (*5*).

For the binding of C3 and histamine to form C3-histamine (C3h), we used an equilibrium dissociation constant of 0.01 M, with an association rate constant ($k_{on}$) of 10 $M^{-1}$ $s^{-1}$ and a dissociation rate constant ($k_{off}$) of 0.1 $s^{-1}$. The spontaneous loss of C3a following C3h formation, resulting in fC3hb generation, was modeled as a first-order reaction with a half-life of 9.6 hours.

In the activation of the complement system by C3h, we assumed the same interactions and parameters used in C-model for C3($H_2O$), except for its inactivation by factor I (FI) in the presence of factor H (FH), where the catalytic constant was reduced to 0.29 $s^{-1}$ to decrease the reaction rate by 4.5-fold compared to the inactivation of C3($H_2O$).

For complement activation by fC3hb, we assumed the same interactions and parameters as those already used in the C-model for fC3b.

> Human C3

MGPTSGPSLLLLLLTHLPLALGSPMYSIITPNILRLESEETMVLEAHDAQGDVPVTVTVHDFPGKKLVLSSEKTVLT
PATNHMGNVTFTIPANREFKSEKGRNKFVTVQATFGTQVVEKVVLVSLQSGYLFIQTDKTIYTPGSTVLYRIFTVNH
KLLPVGRTVMVNIENPEGIPVKQDSLSSQNQLGVLPLSWDIPELVNMGQWKIRAYYENSPQQVFSTEFEVKEYVLPS
FEVIVEPTEKFYYIYNEKGLEVTITARFLYGKKVEGTAFVIFGIQDGEQRISLPESLKRIPIEDGSGEVVLSRKVLL
DGVQNPRAEDLVGKSLYVSATVILHSGSDMVQAERSGIPIVTSPYQIHFTKTPKYFKPGMPFDLMVFVTNPDGSPAY
RVPVAVQGEDTVQSLTQGDGVAKLSINTHPSQKPLSITVRTKKQELSEAEQATRTMQALPYSTVGNSNNYLHLSVLR
TELRPGETLNVNFLLRMDRAHEAKIRYYTYLIMNKGRLLKAGRQVREPGQDLVVLPLSITTDFIPSFRLVAYYTLIG
ASGQREVVADSVWVDVKDSCVGSLVVKSGQSEDRQPVPGQQMTLKIEGDHGARVVLVAVDKGVFVLNKKNKLTQSKI
WDVVEKADIGCTPGSGKDYAGVFSDAGLTFTSSSGQQTAQRAELQCPQPAARRRRSVQLTEKRMDKVGKYPKELRKC
CEDGMRENPMRFSCQRRTRFISLGEACKKVFLDCCNYITELRRQHARASHLGLARSNLDEDIIAEENIVSRSEFPES
WLWNVEDLKEPPKNGISTKLMNIFLKDSITTWEILAVSMSDKKGICVADPFEVTVMQDFFIDLRLPYSVVRNEQVEI
RAVLYNYRQNQELKVRVELLHNPAFCSLATTKRRHQQTVTIPPKSSLSVPYVIVPLKTGLQEVEVKAAVYHHFISDG
VRKSLKVVPEGIRMNKTVAVRTLDPERLGREGVQKEDIPPADLSDQVPDTESETRILLQGTPVAQMTEDAVDAERLK
HLIVTPSG**CGEQ**NMIGMTPTVIAVHYLDETEQWEKFGLEKRQGALELIKKGYTQQLAFRQPSSAFAAFVKRAPSTWL
TAYVVKVFSLAVNLIAIDSQVLCGAVKWLILEKQKPDGVFQEDAPVIHQEMIGGLRNNNEKDMALTAFVLISLQEAK
DICEEQVNSLPGSITKAGDFLEANYMNLQRSYTVAIAGYALAQMGRLKGPLLNKFLTTAKDKNRWEDPGKQLYNVEA
TSYALLALLQLKDFDFVPPVVRWLNEQRYYGGGYGSTQATFMVFQALAQYQKDAPDHQELNLDVSLQLPSRSSKITH
RIHWESASLLRSEETKENEGFTVTAEGKGQGTLSVVTMYHAKAKDQLTCNKFDLKVTIKPAPETEKRPQDAKNTMIL
EICTRYRGDQDATMSILDISMMTGFAPDTDDLKQLANGVDRYISKYELDKAFSDRNTLIIYLDKVSHSEDDCLAFKV
HQYFNVELIQPGAVKVYAYYNLEESCTRFYHPEKEDGKLNKLCRDELCRCAEENCFIQKSDDKVTLEERLDKACEPG
VDYVYKTRLVKVQLSNDFDEYIMAIEQTIKSGSDEVQVGQQRTFISPIKCREALKLEEKKHYLMWGLSSDFWGEKPN
LSYIIGKDTWVEHWPEEDECQDEENQKQCQDLGAFTESMVVFGCPN

| Mass (Da) | Positions | PTMs | Sequence |
|---|---|---|---|
| 3972.841 | 1002-1036 | 0 | HLIVTPSG**CGEQ**NMIGMTPTVIAVHYL:ETEQWEK (unmodified) |
| 4065.910 | 1002-1036 | 1 | HLIVTPSG**CGEQ(Histamine)**NMIGMTPTVIAVHYLDETEQWEK |

**Fig. S1. Predicted peptide mass before and after histamine reaction.**

FASTA-format sequence of full-length human C3, retrieved from UniProt entry CO3_HUMAN at https://www.uniprot.org/uniprotkb/P01024/entry, with the thioester bond residues Cys1010 to Gln1013 highlighted in yellow and bold face, and the expected monoisotopic masses of the containing peptides (amino acids 1002–1036) after free cysteine thiol methylation and complete trypsin digestion of C3 (unmodified) and C3h (histamine adduct). Peptide masses were calculated with EXPASY PeptideMass (https://web.expasy.org/peptide_mass/), selecting Trypsin as the enzyme and allowing for 0 missed cuts.

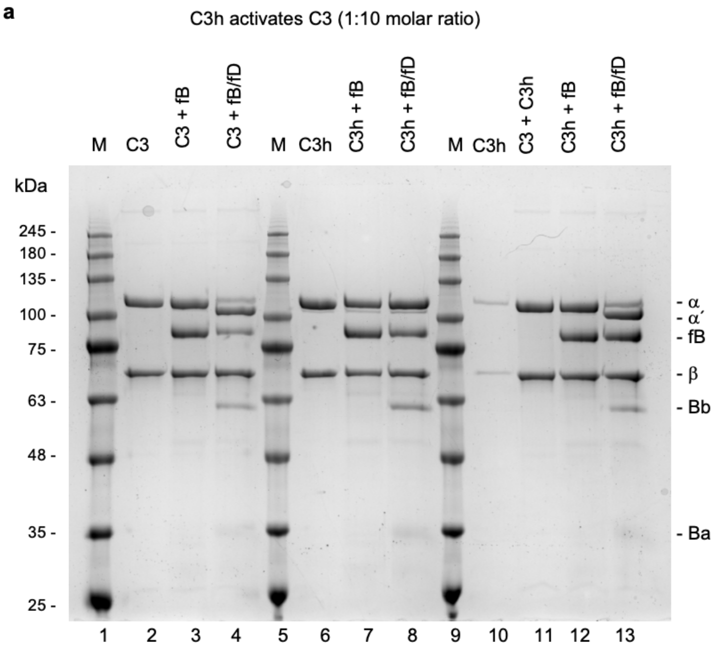


**Fig. S2. C3h forms a C3 convertase downregulated by FH and FI.**

(**A**) Comparison by Coomassie-stained SDS-PAGE gel electrophoresis of the cleavage of purified C3 and C3h by FB and FD (C3, lanes 1-4; C3h, lanes 5-8) with that of C3 supplemented with a 1:10 molar ratio of C3h lanes 9-13). While C3 is rapidly cleaved into C3b and C3a presumably by trace amounts of C3(H2O) convertases, C3h remains intact. By adding a small amount of C3h to C3, the C3 cleavage reaction became accelerated by a factor of ~2 (Fig. 2 in the main text).

a

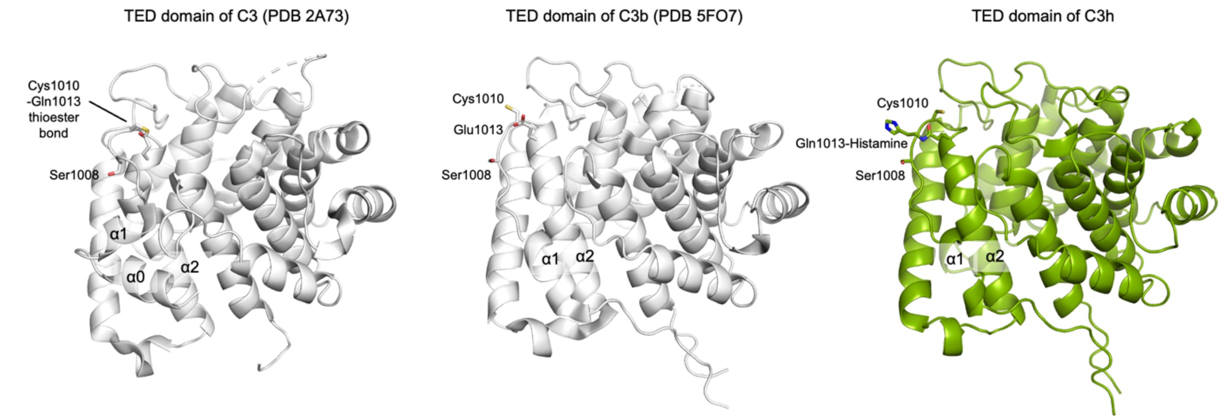


**Fig. S3. The TED domain of C3h has a fully activated conformation.**

Cartoon representation of structurally aligned TED domains from C3 (PDB 2A73), C3b (PDB 5FO7), and C3h (this work). The TED domain of C3h has transitioned toward a fully activated conformation, defined by that of C3b. In particular, the α0-α1 configuration seen in the TED domain of C3 is absent from C3h. The RMSD after superposition between the TED domains of C3b and C3h is 0.68 Å (296 residues) versus 3.04 Å (280 residues) between the TED domains of C3 and C3h.

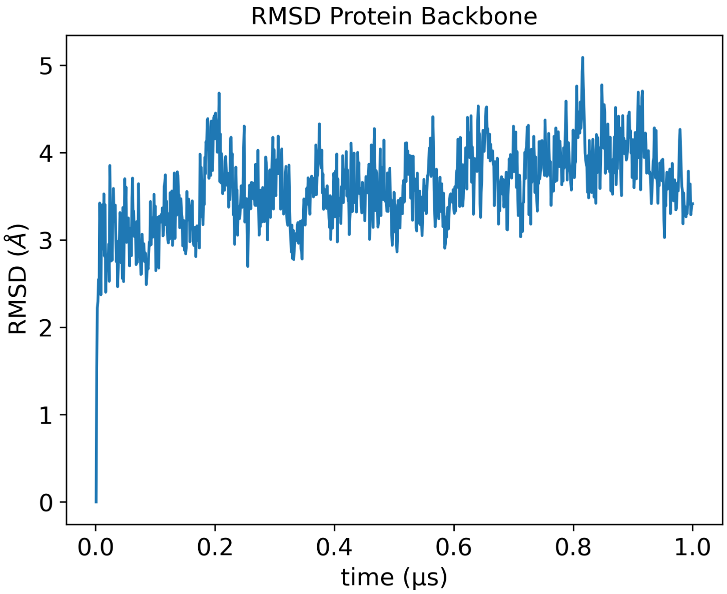


**Fig. S4. Simulations of a C3-like conformation containing the adduct formed after reaction of the thioester bond with histamine.**

The structure was prepared by overlapping the TED domain of C3h, including the histamine covalent adduct, with that of C3 (PDB 2A73) and simulated for 1 μs. The RMSD of the backbone protein atoms shows a stable behavior after ~300 ps.

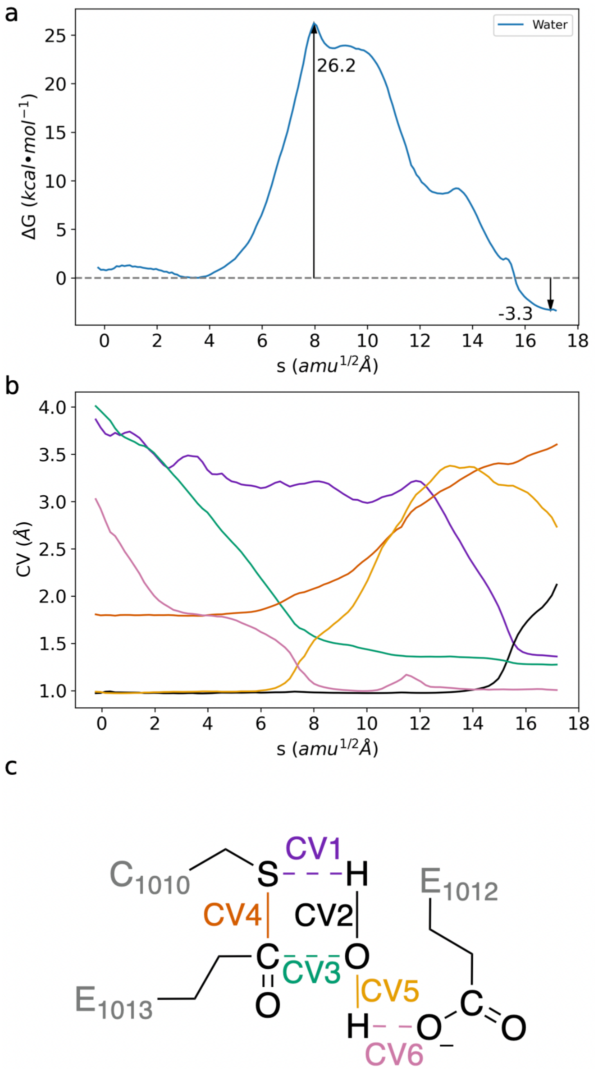


**Fig. S5. Minimum Free Energy Path (MFEP) calculation for the hydrolysis reaction of the thioester bond in C3.**

(**A**) Free energy profile for the hydrolysis reaction. The activation and reaction energies are given in the figure. The structures are shown in Figure 7 of the main text. (**B**) Evolution of the most important distances involved in the reaction along the MFEP. The color code corresponds to the Collective Variables (CVs) definition in panel (**C**). (**C**) Definition of the CVs employed in the determination of the MFEP for the reaction of water with the thioester complex. The QM region employed in the QM/MM simulations includes the water molecule and the side chains of residues C1010, E1012, and E1013.

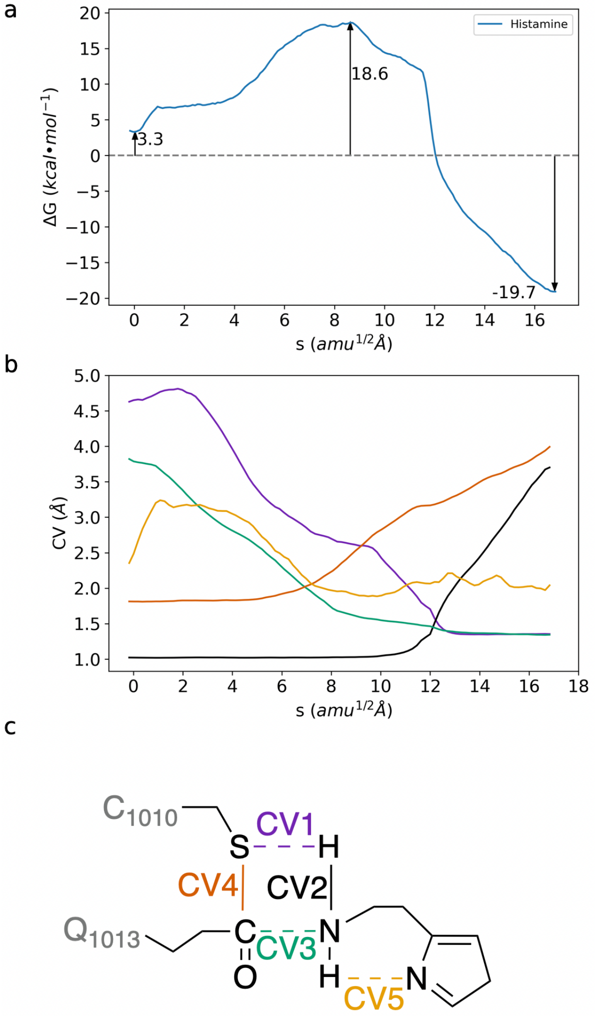


**Fig. S6. Minimum Free Energy Path (MFEP) calculation for the aminolysis reaction of the thioester bond in C3.**

(**A**) Free energy profile for the aminolysis reaction with histamine. The activation and reaction energies are given in the figure. The structures are shown in Figure 7 of the main text. (**B**) Evolution of the most important distances involved in the reaction along the MFEP. The color code corresponds to the Collective Variables (CVs) definition in panel (**C**). (**C**) Definition of the CVs employed in the determination of the MFEP for the reaction of histamine with the thioester complex. The QM region utilized in the QM/MM simulations includes the histamine molecule and the side chains of residues C1010 and Q1013.

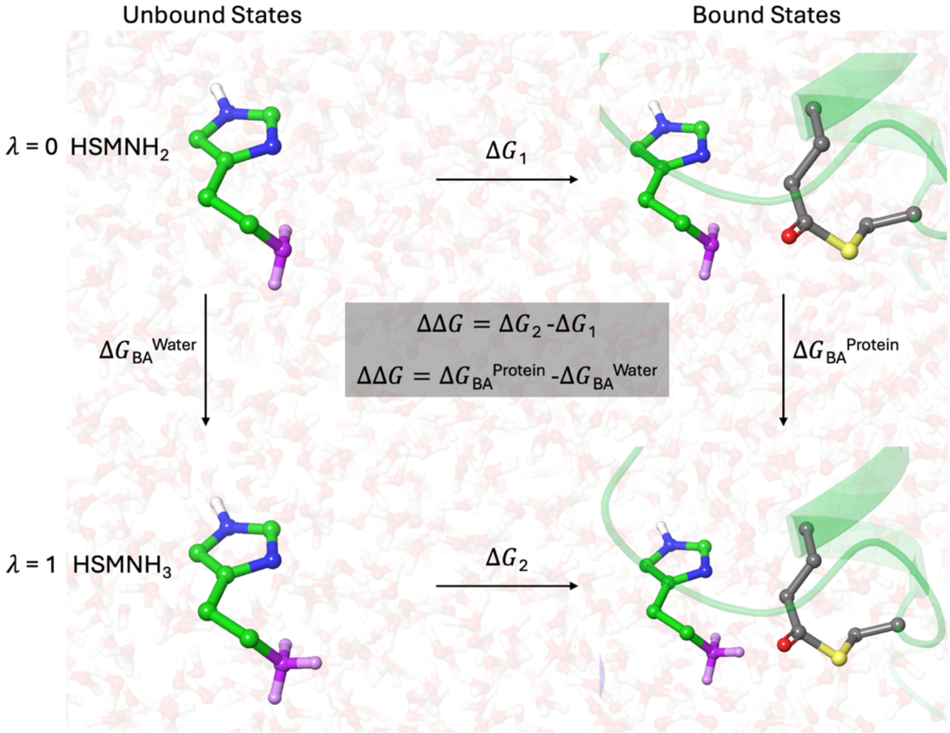


**Fig. S7. Thermodynamic cycle employed to calculate the differences in the binding free energies of protonated and free base forms of the terminal amino group of histamine.**

| (a) Sample details and set-up | |
|---|---|
| Organism | Human |
| Source (Catalog No. or reference) | Natural source with chemical modification as described in the main text |
| Buffer | 20 mM Tris-HCl (pH 8.0), 150 mM NaCl, 3% (w/v) glycerol |
| SAXS data collection mode | Size exclusion chromatography |
| Sample temperature (°C) | 15.0 |
| Sample concentration (mg/ml) | 1.2 |
| Sample injection volume (μl) | 50 |
| SEC column type | Shodex KW-403 (separation range between 10-700 kDa) |
| SEC flow rate (ml/min) | 0.16 |
| **(b) SAS data collection** | |
| Data acquisition/reduction software | SASFLOW / Chromixs |
| Source/instrument description | Diamond B21, Eiger 4M |
| Measured $q$-range ($q_{min} - q_{max}$; $nm^{-1}$) | 0.0045 to 0.1660 |
| Exposure time (s) | ~3 s (12 frames averaged, ~36 s) |
| **(c) SAS-derived structural parameters (from BioXTAS RAW and ATSAS v3.2)** | |
| *Guinier Analysis* | |
| $R_g \pm \sigma$ (Å) | 48.19 ± 0.08 |
| Data point range ($q$ range; $nm^{-1}$) | 0–172 (0.0045–0.02697) |
| Linear fit assessment (fidelity) | 0.9950 |
| *PDDF[1]/P(r) analysis* | |
| $D_{max}$ (Å) | 145 |
| $P(r)$ fit assessment (fidelity) | 0.8759 |
| *Molecular weight estimates (kDa)* | |
| From chemical composition | ~184,000 |
| From SAS (Bayes method) | 169,625 (CI = 151,450–194,950; CI probability = 0.9343) |
| Ab initio *modeling* | |
| Software | *DAMMIF* v3.1.3 (r14636) |
| Envelope volume ($nm^3$) | 438.7 |
| NSD[2] ± $\sigma$ (Å) | 0.9 ± 0.2 |
| Ensemble resolution[3] (Å) | 30.7 |
| *Atomistic modeling methods* | |
| Software | SASREF |
| $\chi^2$ | 1.2 |

**Table S1. SAXS sample details, data collection, analysis, and 3D modeling details.**

1PDDF, Pair distance distribution function.
2NSD, Normalized spatial discrepancy.
3From SASREF.

| HSM-$NH_2$ to HSM-$NH_3^+$ | | Final result: ΔΔG = 0.2 ± 0.6 kcal/mol | | | |
|---|---|---|---|---|---|
| Unbound states | | | Bound states | | |
| System | Replica | ΔG (kcal/mol) | System | Replica | ΔG (kcal/mol) |
| Water | 1 | –44.78 | Protein | 1 | –43.68 |
| Water | 2 | –44.83 | Protein | 2 | –44.17 |
| Water | 3 | –44.91 | Protein | 3 | –45.08 |
| Water | 4 | –44.81 | Protein | 4 | –45.22 |
| Water | 5 | –44.82 | Protein | 5 | –44.76 |
| | Mean | –44.83 | | Mean | –44.6 |
| | Std. dev. | 0.05 | | Std. dev. | 0.6 |

**Table S2. Calculation of the binding free energy difference between the protonated and free base forms of histamine.**

The average result is obtained after five different replicas for the transformation of the free base form into the protonated form in both environments.

### References


1. J. H. Butterfield, A. Ravi, T. Pongdee, Mast Cell Mediators of Significance in Clinical Practice in Mastocytosis. *Immunol. Allergy Clin. North Am.* **38**, 397–410 (2018).

2. W. Lorenz, A. Doenicke, R. Meyer, H. J. Reimann, J. Kusche, H. Barth, H. Geesing, M. Hutzel, B. Weissenbacher, An improved method for the determination of histamine release in man: Its application in studies with propanidid and thiopentone. *Eur. J. Pharmacol.* **19**, 180–190 (1972).

3. T. Boehm, B. Reiter, R. Ristl, K. Petroczi, W. Sperr, T. Stimpfl, P. Valent, B. Jilma, Massive release of the histamine-degrading enzyme diamine oxidase during severe anaphylaxis in mastocytosis patients. *Allergy* **74**, 583–593 (2019).

4. T. Boehm, M. Karer, P. Matzneller, N. Buchtele, F. Ratzinger, K. Petroczi, C. Schoergenhofer, M. Schwameis, H. Burgmann, M. Zeitlinger, B. Jilma, Human diamine oxidase is readily released from activated neutrophils ex vivo and in vivo but is rarely elevated in bacteremic patients. *Int. J. Immunopathol. Pharmacol.* **34**, 2058738420954945 (2020).

5. B. O. Elmore, J. A. Bollinger, D. M. Dooley, Human kidney diamine oxidase: heterologous expression, purification, and characterization. *JBIC J. Biol. Inorg. Chem.* **7**, 565–579 (2002).
